# Fasting-sensitive SUMO-switch on Prox1 controls hepatic cholesterol metabolism

**DOI:** 10.1101/2022.08.17.504229

**Authors:** Ana Jimena Alfaro Nunez, Claudia Dittner, Janina Becker, Anne Loft, Amit Mhamane, Adriano Maida, Anastasia Georgiadi, Phivos Tsokanos, Katarina Klepac, Eveline Molocea, Rabih Merahbi, Karsten Motzler, Julia Geppert, Rhoda Anane Karikari, Julia Szendrödi, Annette Feuchtinger, Susanna Hofmann, Frauke Melchior, Stephan Herzig

## Abstract

The liver is the major metabolic hub, ensuring appropriate nutrient supply during fasting and feeding. In obesity, accumulation of excess nutrients hampers proper liver function and is linked to non-alcoholic fatty liver disease. Understanding the signaling mechanisms that enable hepatocytes to quickly adapt to dietary cues, might help to restore balance in liver diseases. Post-translational modification by attachment of the Small Ubiquitin-like Modifier (SUMO), allows for a dynamic regulation of numerous processes including transcriptional reprograming. Here, we demonstrate that the specific SUMOylation of transcription factor Prox1 represents a nutrient-sensitive determinant of hepatic fasting metabolism. Prox1 was highly modified by SUMOylation on lysine 556 in the liver of ad libitum and re-fed mice, while this modification was strongly abolished upon fasting. In a context of diet-induced obesity, Prox1 SUMOylation became insensitive to fasting cues. Hepatocyte-selective knock in of a SUMOylation-deficient Prox1 mutant into mice fed a high fat/high fructose diet led to reduction of systemic cholesterol levels, associated with the induction of bile acid detoxifying pathways in mutant livers during fasting. As appropriate and controlled fasting protocols have been shown to exert beneficial effects on human health, tools to maintain the nutrient-sensitive SUMOylation switch on Prox1 may thus contribute to the development of “fasting-based” approaches for the maintenance of metabolic health.

## Introduction

Hepatic metabolism critically regulates systemic energy homeostasis, in part by controlling the adaptation response during variations in nutrient availability and detoxification processes. Hepatocytes have evolved to sense and react to dietary signals through a network of nutrient sensing and signal transduction pathways in order to cope with the fluctuating energy demands (Bechmann et al., 2012; Juza & Pauli, 2014; Panda et al., 2002; Si-Tayeb et al., 2010).

Physiological cues such as fasting and feeding signals are translated into specific transcriptional metabolic programs, which enable long-term regulation and adaptation of cellular metabolism. The activity of key regulatory factors, including transcription factors, is modulated by reversible post-translational protein modifications, which allows for fast adaptive responses to changes in the cellular environment. The regulatory network driven by post-translational modifications in response to metabolic cues has mainly been studied in the context of phosphorylation. However, one of the post-translational modifications that is gaining attention as a key regulator of transcription is SUMOylation.

SUMOylation involves the covalent attachment of the 11 kDa Small Ubiquitin-like Modifier (SUMO) with the help of E1, E2 and E3 enzymes; SUMOylation is a reversible and very dynamic modification due to the highly active isopeptidases that catalyze SUMO de-conjugation (Matunis et al., 1996) (Geiss-Friedlander & Melchior, 2007; Mahajan et al., 1997). The conjugation of SUMO occurs at lysine residues frequently within a SUMOylation consensus motif (YKxE), a recognition site for the E1/E2 SUMO loading complex (Flotho & Melchior, 2013; Gareau & Lima, 2010). Mammals express two SUMO families, SUMO1 and SUMO2/3. Although conjugated via the same enzymatic pathway, the SUMO1 and SUMO2/3 isoforms share only about 50 % of their amino acid sequence. Consistent with the differences observed between the amino acid sequences, SUMO1 and SUMO2/3 have distinct mechanistic functions and exert different consequences on the target protein (Alegre & Reverter, 2011; Chang et al., 2011; Di Bacco et al., 2006; Tatham et al., 2001; Zhu et al., 2009). Similar to phosphorylation, SUMOylation serves as a molecular switch controlling an array of cellular processes. The attachment of SUMO can generate new binding interfaces or allosterically blocking a binding site thereby modulating the function of the target protein (Flotho & Melchior, 2013; Geiss-Friedlander & Melchior, 2007; Vertegaal, 2022). By promoting alterations in nuclear localization, preventing protein degradation, changing DNA binding affinity and regulating the interaction with chromatin modifying complexes, SUMOylation controls the activity of transcription factors and co-regulators thus playing a crucial role in the dynamic regulation of transcription (Rosonina, 2019; Treuter & Venteclef, 2011).

Recent developments in SUMO-proteome analysis led to identification of several thousand proteins that are under control of SUMOylation, including a vast number of transcription factors and transcriptional regulators (Barysch et al., 2014; Becker et al., 2013; Correa-Vázquez et al., 2021). Previous studies have demonstrated that SUMOylation participates in the regulation of liver metabolism by controlling the activity of key hepatic transcription factors (Balasubramaniyan et al., 2013; Kim et al., 2015; Lee et al., 2014; Stein et al., 2014).

Thus, we aimed to identify the dynamic role of SUMOylation in the transcriptional control of liver metabolism in response to changes in nutrient availability. To this end, we performed a SUMO-proteome analysis of the mouse liver in the fasted and re-feeding cycle. We identified the fasting-sensitive SUMOylation of the transcription factor Prox1 as a major SUMOylation event in the liver and as a key determinant of systemic cholesterol metabolism. The SUMO- switch on Prox1 allows the regulation of a distinct subset of genes involved in the hepatic cholesterol detoxification system in response to fasting. Our study provides the first example of a fasting-feeding sensitive post-translational modification with immediate functional impact on physiological responses to changes in environmental conditions.

## Experimental Procedures

### Animal studies

The Prox1_K556R_ K.I. line generated by Taconic Biosciences was backcrossed with wild type C57BL/6N mice and expanded in the animal facility of Helmholtz Munich. 8 weeks-old Prox1_K556R_ K.I. (f/f) mice were injected with an AAV_LP1_Cre_wt_ (Cre_AAV) to induce the expression of a K556R Prox1 mutant in hepatocytes. An AAV_LP1_Cre_mut_ (Ctrl_AAV) coding for an untranslatable Cre was used as control. The generation of the AAV constructs and the control for hepatocyte specificity is described in detail in the supplemental experimental procedures.

Prox1_K556R_ K.I. and wild type C57BL/6N (Charles River laboratories) were maintained on a 12/12 fh light/dark cycle and were fed a regular chow diet ad libitum unless indicated otherwise. For the metabolic challenges mice were fed either a 60 % high fat diet (Research Diets - D12492i) or a combination of 45 % high fat diet (Research Diets - D12451i) and fructose water (20% w/v).

Fasting and re-feeding schedules are explained in the main text and in the figure legends; Zeitgeber (ZT) 0 = lights on and ZT12 = lights off.

Protocols for in vivo characterization and liver lipid analysis are explained in the supplemental experimental procedures. All animal studies were performed in accordance with German animal welfare legislation and in specific pathogen-free conditions in the animal facility of the Helmholtz Center, Munich, Germany. Protocols were approved by the Institutional Animal Welfare Officer (Tierschutzbeauftragter), and necessary licenses were obtained from the state ethics committee and government of Upper Bavaria (ROB-55.2-2532.Vet_02-17-49 and ROB- 55.2-2532.Vet_02-15-164).

### Tissue collection, serum analysis and lipoprotein profile by fast protein liquid chromatography

The body weight and the blood glucose levels were recorded in every study prior to tissue collection. Mice were sacrificed by cervical dislocation and decapitated immediately. The blood was collected in serum gel tubes, incubated at room temperature for 5-10 min and store at 4° C until further processing. The collection of liver samples from wild type mice was done with a freeze clamping technique where the liver was exposed and pre-frozen (liquid nitrogen) forceps were used to collect a piece of the left lobe. For the characterization of Prox1_K556R_ K.I. mice the tissues were collected, weighted, washed in PBS and snap-frozen in liquid nitrogen. A sample from the medial liver lobe was placed on a histocasette and incubated in formalin for fixation. Once all tissues were collected the blood samples were centrifuged at 2000 g for 10 min at 4° C; the serum was collected and snap-frozen in liquid nitrogen. Tissue and serum samples were stored at -80° C.

Individual serum samples were analyzed using an automatized system (Beckman Coulter AU480 Chemistry Analyzer).

To generate a lipoprotein profile serum samples were pooled and subjected to gel filtration by fast protein liquid chromatography using a Superose 6 10/300 GL column (Cytiva 17-5172-01). The cholesterol content of individual fractions was determined using a total cholesterol assay kit (Invitrogen A12216).

### Immunoblot analysis and Immunoprecipitation

Around 20 mg of frozen liver tissue were lysed and homogenized using a tissue lyser (Retsch - MM400) with steel beads in 500 μl ice cold lysis buffer containing Tris (50 mM) pH 6.8, EDTA (1 mM), NaCl (150 mM), Igepal (1 %) and N-Ethylmaleimide (100 mM) as isopeptidase inhibitor (Sigma - E3876) supplemented with protease and phosphatase inhibitors in tablets (Roche - 42484600 and 4906845001 respectively). Liver lysates were analyzed by immunoblotting using anti-Prox1 (Millipore - 07-537 and Abcam - ab38692 in Fig. 3B), anti-Phospho_S6K_(Thr389)_ (Cell Signaling - 9234), anti-Cre Recombinase (Cell Signaling - 15036), goat anti-RanGAP1 (produced by our collaborator at the Melchior lab), anti-β Actin (Sigma - A5441) and anti-Vcp (Abcam - ab11433) antibodies. Blots were analyzed and quantified with Image Lab (Bio-Rad Laboratories) SUMO immunoprecipitation from liver tissue samples has was performed as described (Barysch et al., 2014; Becker, 2012; Becker et al., 2013).

### Gene expression analysis

Around 5 mg of liver tissue were homogenized using a tissue lyser (Retsch - MM400) with steel beads in 1 ml Trizol (Thermo Scientific - 15596018). The samples were mixed with 0.2 ml chloroform, the aqueous phase was collected and mixed with 0.6x volumes of 100 % ethanol. The samples were loaded on a EconoSpin column and the RNA was washed 3 times with RPE buffer (QIAGEN). The RNA was eluted with distilled nuclease free water. The RNA concentration was determined with a NanoDrop 2000 spectrophotometer (Thermo Scientific) and the integrity was controlled by gel electrophoresis using a RNA 6000 Nano Kit (Agilent) according to the manufacturer’s protocol using a 2100 Bioanalyzer (Agilent). RNA samples were diluted to a 100 ng/μl concentration and stored at -80° C.

For the qPCR expression analysis, cDNA was generated from 1 μg of RNA with the QuantiTect Reverse Transcription Kit (QIAGEN) according to the manufacturer’s protocol. The expression of selected genes was analyzed using the TaqMan Gene Expression reagents (Thermo Scientific) with the QuantStudio 6 Flex Real-Time PCR system (Thermo Scientific).

The RNA sequencing libraries were generated by Novogene Europe. The adapter sequence from the raw files was removed by using Cutadapt 4.1. Raw counts were then aligned to the mouse reference genome using STAR 2.7.10a. Any genes that have no transcript detected in any samples were removed. Data normalization and differential expression analysis was performed using the DESeq2 R-package from Bioconductor (Love, Huber, and Anders, 2014). A statistical threshold was set to an adjusted pvalue (padj) < 0.05; DESeq2 uses the Benjamini-Hochberg to adjust for multiple testing. A threshold for the effect size was set to a log2 fold change (FC) of < -0.5 or > 0.5. Analysis of the differentially expressed genes for biological pathways was performed using the enrichKEGG function (Kyoto Encyclopedia of Genes and Genomes database).

### Overexpression of HA-Prox1 constructs and invitro SUMOylation assay

The HA-Prox1 wild type, K556R and E558A mutant constructs were generated as described (Dittner, 2016). HepG2 cells were transfected using Lipofectamine LTX with Plus Reagent (Thermo Fisher Scientific) according to the manufacturers protocol. HepG2 cell lysates were analyzed by immunoblotting using anti-HA (Covance - MMS-101P) antibodies.

In vitro SUMOylation reactions were done with recombinant E1 Aos1/Uba2 (100 nM) and E2 Ubc9 (200 nM) in the presence of either SUMO1 or SUMO2 (5 μM). The reactions were carried out in a volume of 20 μl assay buffer: HEPES/KOH (20mM) pH 7.3, KAcO (110 mM), Mg(AcO)_2_ (2 mM), EGTA (1 mM), DTT (1 mM) and Tween 20 (0.05 % v/v) supplemented with protease inhibitors and ovalbumin (0.2 mg/ml). The reactions were incubated with purified mouse wild type or K556R Prox1 at 30° C. Reactions were initiated by adding ATP (1 mM) and stopped by the addition of 20 μl 2x SDS sample buffer. Purification of mouse Prox1 is described in detail in the supplemental experimental procedures. Recombinant E1, E2 and SUMO proteins were purified as described previously (Pichler et al., 2004; Werner et al., 2009)

### Statistical Analyses

Data expressed as means ± SEM. Comparison of time-course experiments was assessed by one-way ANOVA with Dunnett’s multiple comparison test relative to samples collected at ZT 12.

Multiple group comparisons were assessed by one-way ANOVA with Tukeýs multiple comparison test between groups. Multiple group comparisons of two factors (fasted and refed) were assessed by two-way ANOVA with Sidak’s multiple comparison test between different groups and conditions. P≤0.05 was considered statistically significant. * P≤0.05, ** P≤0.01, *** P≤0.001, **** P≤0.0001.

## Results

### Prox1 is modified by SUMO at lysine 556 in response to nutrient availability

In order to define the hepatic SUMO-proteome in response to changes in nutrient availability, we extracted endogenous SUMO targets from liver tissue of fasted (16 h) and re-fed (fasted 16 h and re-fed 2 h) wild-type mice using monoclonal anti-SUMO1 and anti-SUMO2/3 antibodies and subsequent peptide elution as described previously (Barysch et al., 2014; Becker et al., 2013). The isolated proteins were analyzed by mass spectrometry (Fig. 1A) and over two hundred SUMO candidates were identified (Becker, 2012). Several candidates were differentially modified between the fasted and re-fed states (Table 1).

**Fig. 1.**
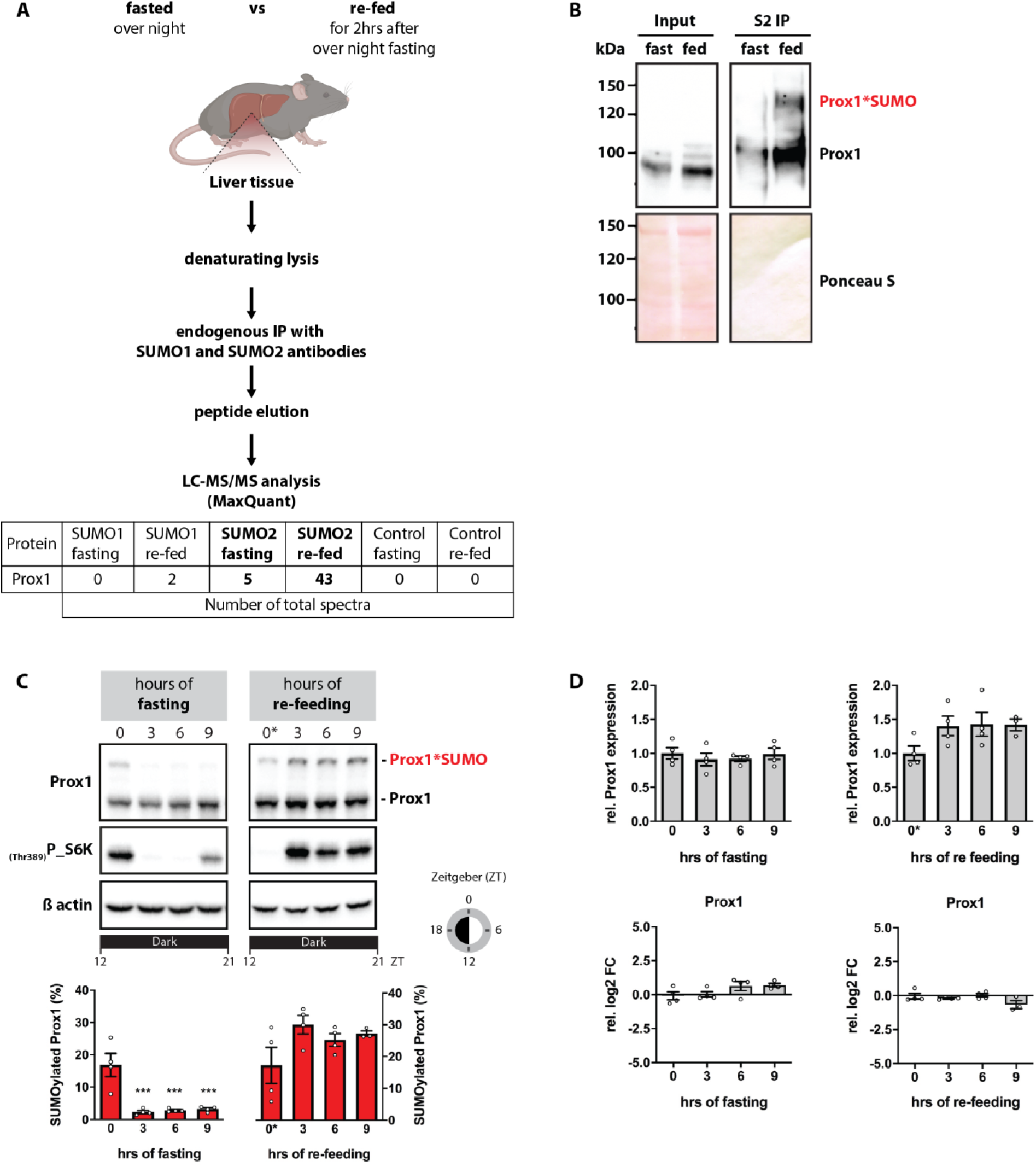
Hepatic Prox1 is modified by SUMOylation in response to in response to nutrient availability. **A)** Schematic representation of the protocol used to enrich and detect endogenous SUMO targets in the mouse liver in the fasted (16 h) and re-fed (fasted 16 h and re-fed 2 h) states (Becker, 2012; Becker et al., 2013). **B)** SUMO2 immunoprecipitation using crosslinked SUMO2 antibody beads (100 μl) and liver lysates (10 mg protein) from wild-type C57BL6/J mice samples, fasted: 16 h or re-fed: 16 h fasted and re-fed for 2 h. Eluates were analyzed by immunoblotting using anti-Prox1 antibodies, ponceau staining was used as loading control. **C)** 8 weeks-old C57BL/6N male mice were fasted: the food was removed at ZT 12 or re-fed: all groups were fasted for 8 h during the light phase (ZT 4-12) for synchronization, the food was re- introduced at ZT 12. Tissue samples were collected at ZT 12, 15, 18 and 21 (n=4). Liver lysates were analyzed by immunoblotting using anti-Prox1 and anti-P_S6K(Thr389) antibodies; β actin was detected for input control. The quantification of SUMOylated Prox1 (%) is shown. **D)** Quantification of total Prox1 protein expression as well as Prox1 mRNA levels analyzed by qPCR are shown. qPCR data presented as relative log2 fold change (FC) normalized to the housekeeping gene TBP. Every dot represents one individual mouse. Data: mean ±SEM. Significance was determined by one-way ANOVA with Dunnett’s multiple comparison test relative to samples collected at ZT 12. * P≤0.05, ** P≤0.01, *** P≤0.001, **** P≤0.0001.

**Table 1.**
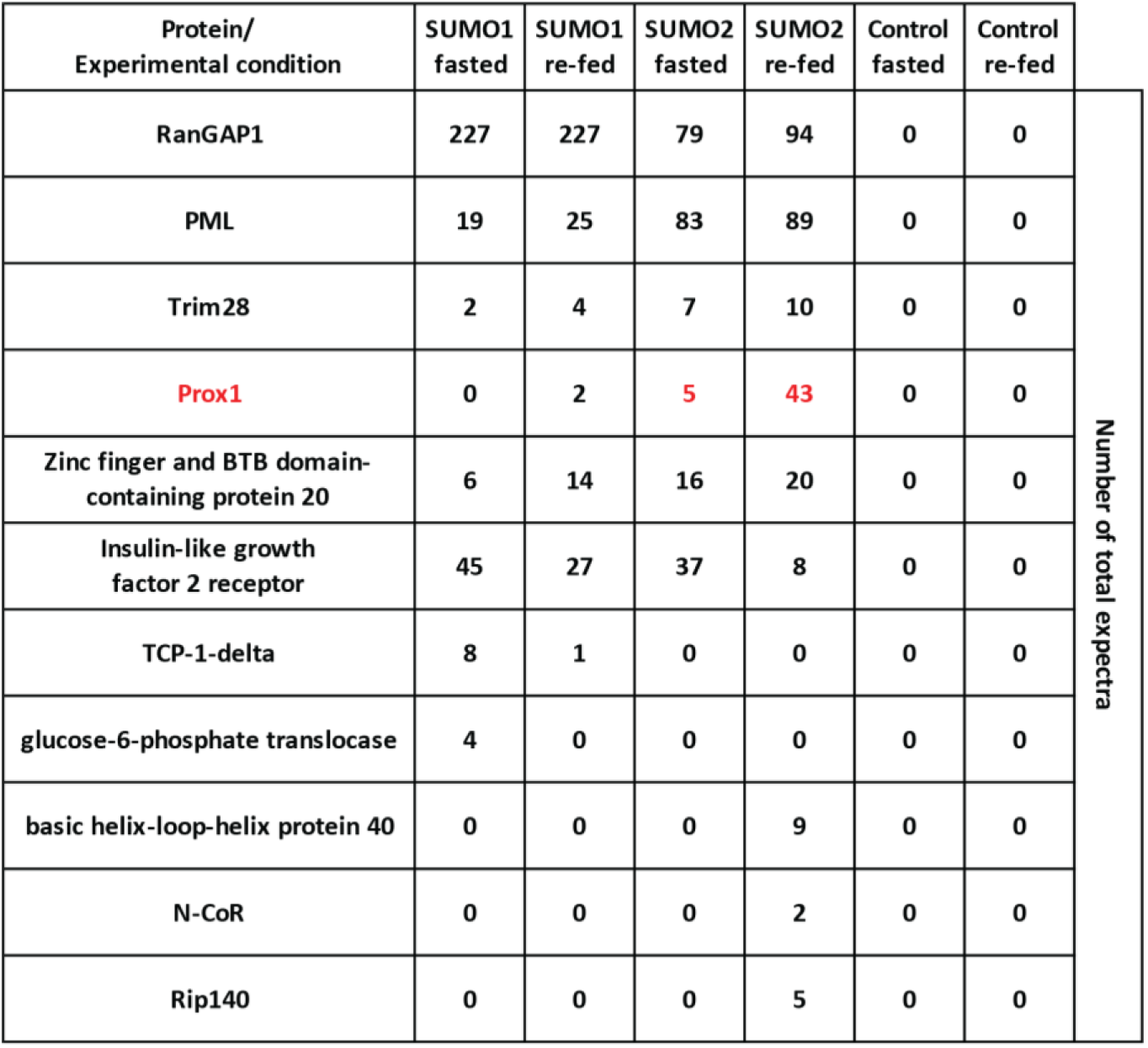
SUMOylated proteins in the mouse liver during fasting or re-fed states. An immunoprecipitation using SUMO1 and SUMO2/3 antibodies was performed using liver tissue of fasted or re-fed mice according to (Barysch et al., 2014; Becker, 2012; Becker et al., 2013). The isolated proteins were analyzed by mass spectrometry. Table showing the peptide count for proteins differentially modified between the fasted and re-fed states (Becker, 2012).

| Protein/<br>Experimental condition | SUMO1<br>fasted | SUMO1<br>re-fed | SUMO2<br>fasted | SUMO2<br>re-fed | Control<br>fasted | Control<br>re-fed | Number of total expectra |
| --- | --- | --- | --- | --- | --- | --- | --- |
| RanGAP1 | 227 | 227 | 79 | 94 | 0 | 0 |  |
| PML | 19 | 25 | 83 | 89 | 0 | 0 |  |
| Trim28 | 2 | 4 | 7 | 10 | 0 | 0 |  |
| Prox1 | 0 | 2 | 5 | 43 | 0 | 0 |  |
| Zinc finger and BTB domain-<br>containing protein 20 | 6 | 14 | 16 | 20 | 0 | 0 |  |
| Insulin-like growth<br>factor 2 receptor | 45 | 27 | 37 | 8 | 0 | 0 |  |
| TCP-1-delta | 8 | 1 | 0 | 0 | 0 | 0 |  |
| glucose-6-phosphate translocase | 4 | 0 | 0 | 0 | 0 | 0 |  |
| basic helix-loop-helix protein 40 | 0 | 0 | 0 | 9 | 0 | 0 |  |
| N-CoR | 0 | 0 | 0 | 2 | 0 | 0 |  |
| Rip140 | 0 | 0 | 0 | 5 | 0 | 0 |  |

The target showing the most notable difference was Prospero homeobox protein 1 (Prox1). Prox1 is a key transcription factor controlling liver development and metabolism. Prox1 expression is essential for hepatoblast migration and hepatocyte cell commitment (Burke & Oliver, 2002; Dudas et al., 2004; Lu et al., 2021; Sosa-Pineda et al., 2000; Velazquez et al., 2021). In the adult liver, Prox1 controls key aspects of lipid metabolism; adult mice lacking Prox1 in hepatocytes show a strong degree of liver steatosis and hepatic injury (Armour et al., 2017; Dittner, 2016; Goto et al., 2017). Several studies have reported that Prox1 acts as a co- repressor of key hepatic nuclear receptors and transcription factors such as the steroid and xenobiotic receptor (SXR), liver receptor homolog-1 (LRH-1), hepatocyte nuclear factor 4α (HNF4α) and estrogen-related receptor α (ERRα) (Armour et al., 2017; Azuma et al., 2011; Charest-Marcotte et al., 2010; Dufour et al., 2011; Qin et al., 2004; Stein et al., 2014). In addition, it has been shown that the transcriptional activity of Prox1 is modulated by the attachment of SUMO1 in endothelial cells (Pan et al., 2009; Shan et al., 2008).

Our mass spectrometry data indicated that Prox1 was substantially SUMOylated in the re-fed state and showed a paralog preference for SUMO2/3 over SUMO1. However, Prox1 SUMOylation was weak during fasting. To confirm these results, we performed a SUMO2 immunoprecipitation using liver tissue from fasted and re-fed wild-type mice. Indeed, the conjugation of Prox1 to SUMO2 was higher in the re-fed state (Fig. 1B). Note, that the SUMOylated species of Prox1 (approx. 120kDA) was easily detected by immunoblotting in liver lysates without prior enrichment (Fig. S1A). To further understand how Prox1 was modified in response to nutrient availability, we analyzed liver samples collected after various fasting or re- feeding time points throughout the dark phase to avoid circadian off-target effects. At the beginning of the dark phase (time point = 0), 15 % of the total Prox1 pool was modified by SUMOylation in ad libitum fed mice. The conjugation of Prox1 with SUMO was lost after 3 h of fasting and remained un-modified for the rest of the dark phase. After re-feeding, the levels of Prox1 SUMOylation were maintained with a tendency to increase over time (Fig. 1C). Prox1 mRNA and protein levels were constant between the fasted and re-fed conditions and through the dark phase (Fig. 1D). Changes in blood glucose and insulin levels as well as the expression of metabolic and circadian genes were measured to corroborate the nutritional state of the mice upon fasting and re-feeding (Fig. S1B). These findings identified a SUMO-switch on Prox1 with a clear response to fasting-feeding signaling.

To gain more insight into the molecular regulation of Prox1 SUMOylation, we aimed to identify the SUMO target site on Prox1. Two SUMOylation consensus motifs on Prox1 have previously been reported around lysine residues 353 and 556 (Shan et al., 2008) (Fig. 2A). We focused on lysine 556; for this, we generated expression constructs coding for mouse wild-type Prox1 (wt), a lysine 556 to arginine (R) mutant (K556R) as well as a glutamic acid 558 to alanine (A) mutant (E558A). The K556R mutant lacks the target lysine while the E558A retains the target lysine but has a mutated SUMOylation consensus motif and should lose the ability to interact with the SUMO loading complex (Flotho & Melchior, 2013; Mahajan et al., 1997; Matunis et al., 1996; Vertegaal, 2022). The status of Prox1 SUMOylation was analyzed using lysates of HepG2 cells over-expressing the Prox1 constructs. Both the un-modified and the SUMOylated species of Prox1 were detected in cells expressing the wt Prox1. However, the conjugation of Prox1 with SUMO was abolished when the target lysine 556 was mutated or the surrounding SUMOylation consensus motif was disrupted (Fig. 2B left), establishing lysine 556 as the main SUMOylation site on Prox1 in a hepatocyte cell model.

**Fig. 2.**
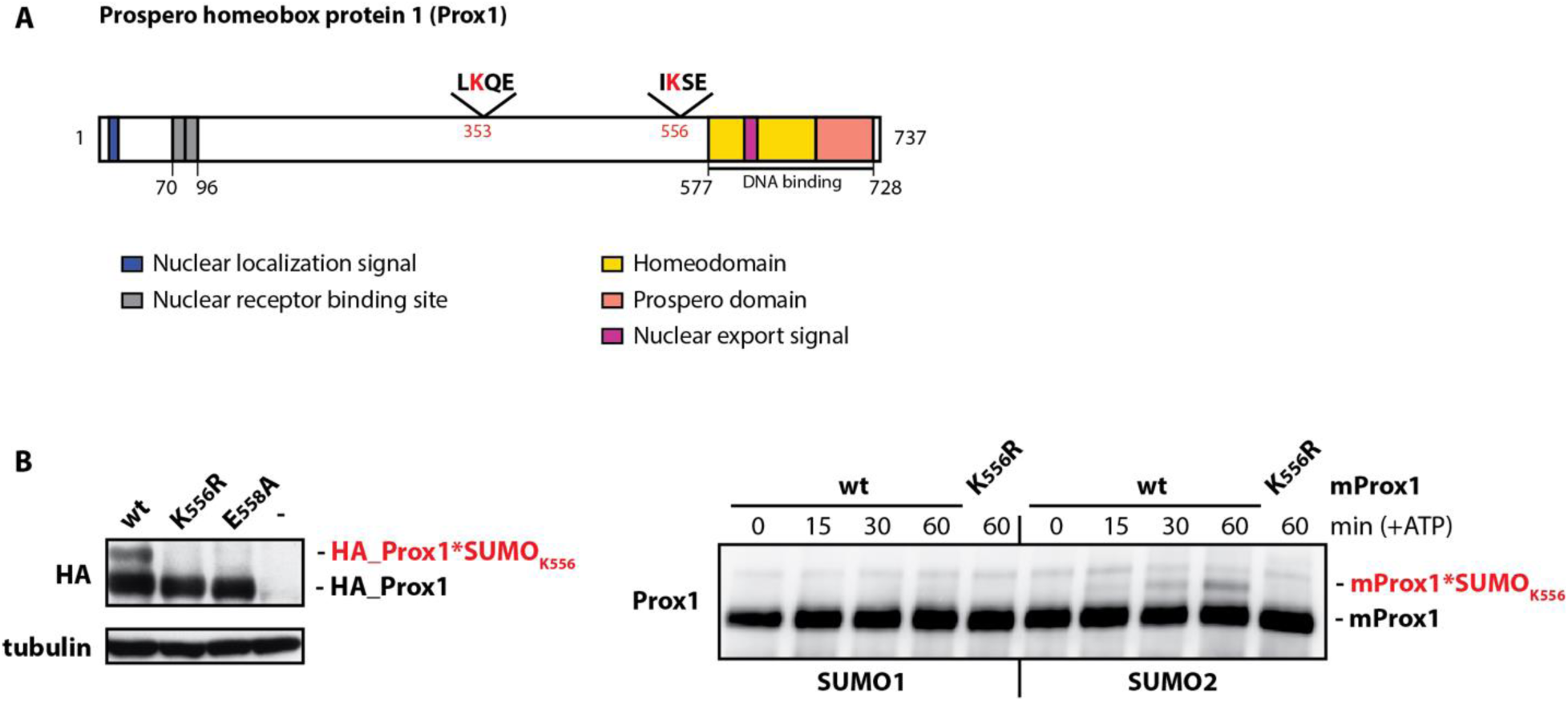
Prox1 is modified by SUMO2 on lysine residue 556. **A)** Schematic representation of Prox1. Functional domains are highlighted. The two putative SUMOylation consensus motifs (ΨKXE) with SUMOylation sites at lysine (K) residues 353 and 556 are marked. Figure adapted from (Elsir et al., 2012). **B) Left:** HepG2 cells were transfected with mouse HA-tagged wild-type Prox1 (wt), K556R mutant Prox1 or E558A mutant Prox1. Lysates were analyzed by immunoblotting using anti-HA antibodies; tubulin was detected for input control. **B) Right:** Mouse un-tagged Prox1 (mProx1) was purified from HEK293T cells. Purified mProx1 (100 nM) was incubated with recombinant E1 enzyme (100 nM) and E2 enzyme (200 nM) together with either SUMO1 or SUMO2 (5 μM) at 30°C. The enzymatic reactions were started with ATP (1 mM) and incubated for 15, 30 and 60 min. A reaction with the K556R mutant (KR) was used as a control. The 0 min sample was incubated without ATP. Samples were analyzed by immunoblotting using anti-Prox1 antibodies.

Furthermore, we considered the potential paralog preference for SUMO2 over SUMO1. For this, un-tagged mouse Prox1 (mProx1) was purified from HEK293T cells and used for in vitro SUMOylation assays. The purified mProx1 was incubated with purified SUMO loading enzymes together with either SUMO1 or SUMO2. The in vitro SUMOylation of Prox1 was more efficient with SUMO2 than with SUMO1 suggesting that the paralog preference for SUMO2 was mediated at the level of conjugation (Fig. 2B right).

Given that Prox1 plays a key role in the regulation of liver lipid metabolism (Armour et al., 2017; Dittner, 2016), we decided to investigate the behavior of the Prox1 SUMO-switch in the liver of mice coping with a metabolic lipid burden. For this, adult wild type male mice were fed with a high fat (60 %) diet (HFD) and a standard chow diet was used as a control. After 8 weeks, the HFD group had gained significantly more weight than the control mice (Fig. 3A left). Liver samples were collected 3 h into the dark phase in the fasted and re-fed states. As observed previously, conjugation of Prox1 with SUMO was absent during fasting and present during re- feeding in the liver of lean mice (Fig. 3A). However, in the liver of HFD-fed obese mice the conjugation of Prox1 with SUMO was only mildly affected during fasting (Fig. 3A). Note that both the un-modified and the SUMOylated species of Prox1 were detected in both states (Fig. 3A). The expression of Prox1 at the mRNA and protein levels were constant between both groups (Fig. 3B). These results suggested that under conditions of diet-induced obesity the SUMO-switch on Prox1 was mostly un-responsive to nutrient availability.

**Fig. 3.**
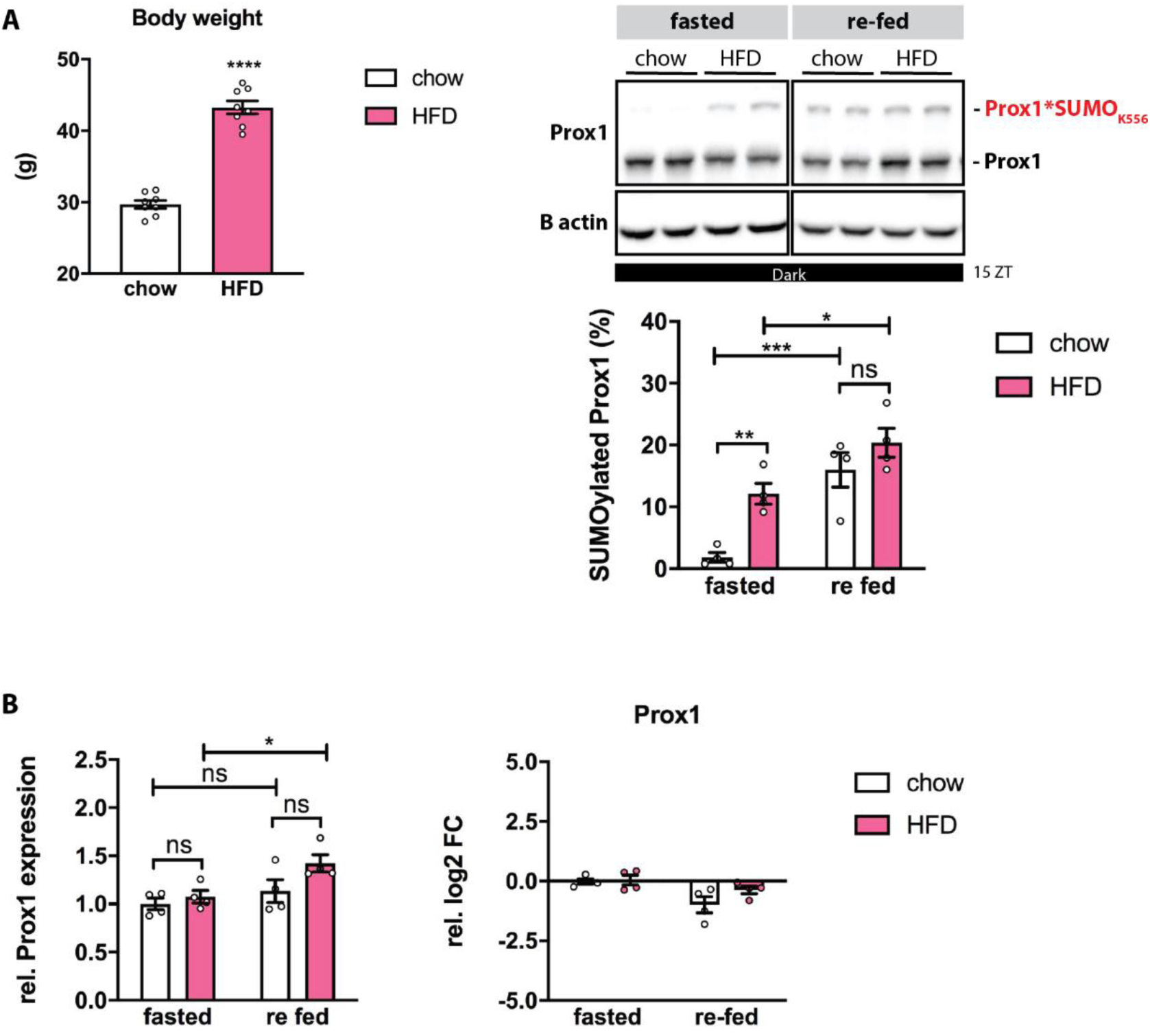
Diet-induced obesity prevents Prox1 de-SUMOylation during fasting. **A)** 6 weeks-old C57BL/6N male mice were fed either a standard chow diet or a high fat (60 %) diet (HFD) for 8 weeks. Left: Body weight records taken a day before tissue collection (n= 8). Fasted for 8 h, the food was removed at ZT 7. Re fed mice were fasted for 8 h (ZT 4-12) for synchronization, the food was re-introduced at ZT 12. Tissues were collected at ZT 15 (n=4). Right: Liver lysates were analyzed by immunoblotting using anti-Prox1 antibodies; β actin was detected for input control. The quantification of SUMOylated Prox1 is shown. **B)** Quantification of total Prox1 protein expression as well as Prox1 mRNA levels analyzed by qPCR are shown. qPCR data presented as relative log2 fold change (FC) normalized to the housekeeping gene TBP. Every dot represents one individual mouse. Body weight data: Data: mean ±SEM. Significance was determined by two-tailed, unpaired t-test. Expression data: mean ±SEM. Significance was determined by two way ANOVA with Sidak’s multiple comparison test between different groups and conditions. *P≤0.05, ** P≤0.01, *** P≤0.001, **** P≤0.0001.

### Liver-specific loss of lysine 556 SUMOylation on Prox1 decreases systemic cholesterol levels

Our previous results raised the question whether the SUMO-switch on Prox1 was required for functional hepatic metabolism.

To this end, we generated a conditional SUMO-deficient Prox1 knock-in mouse model (Prox1_K556R_ K.I. mice). Upon Cre-recombination the genomic region coding for the wild-type variant was removed allowing for the expression of a Prox1 K556R mutant (Fig. 4A). Using an adeno-associated virus (AAV) as a vector to overexpress the Cre recombinase protein under the control of the hepatocyte-specific LP1 promoter (Cre_AAV), we were able to induce Cre-recombination only in hepatocytes and only after the liver had fully developed. An AAV vector coding for an untranslatable Cre recombinase under the LP1 promoter was used as a control (Ctrl_AAV).

**Fig. 4.**
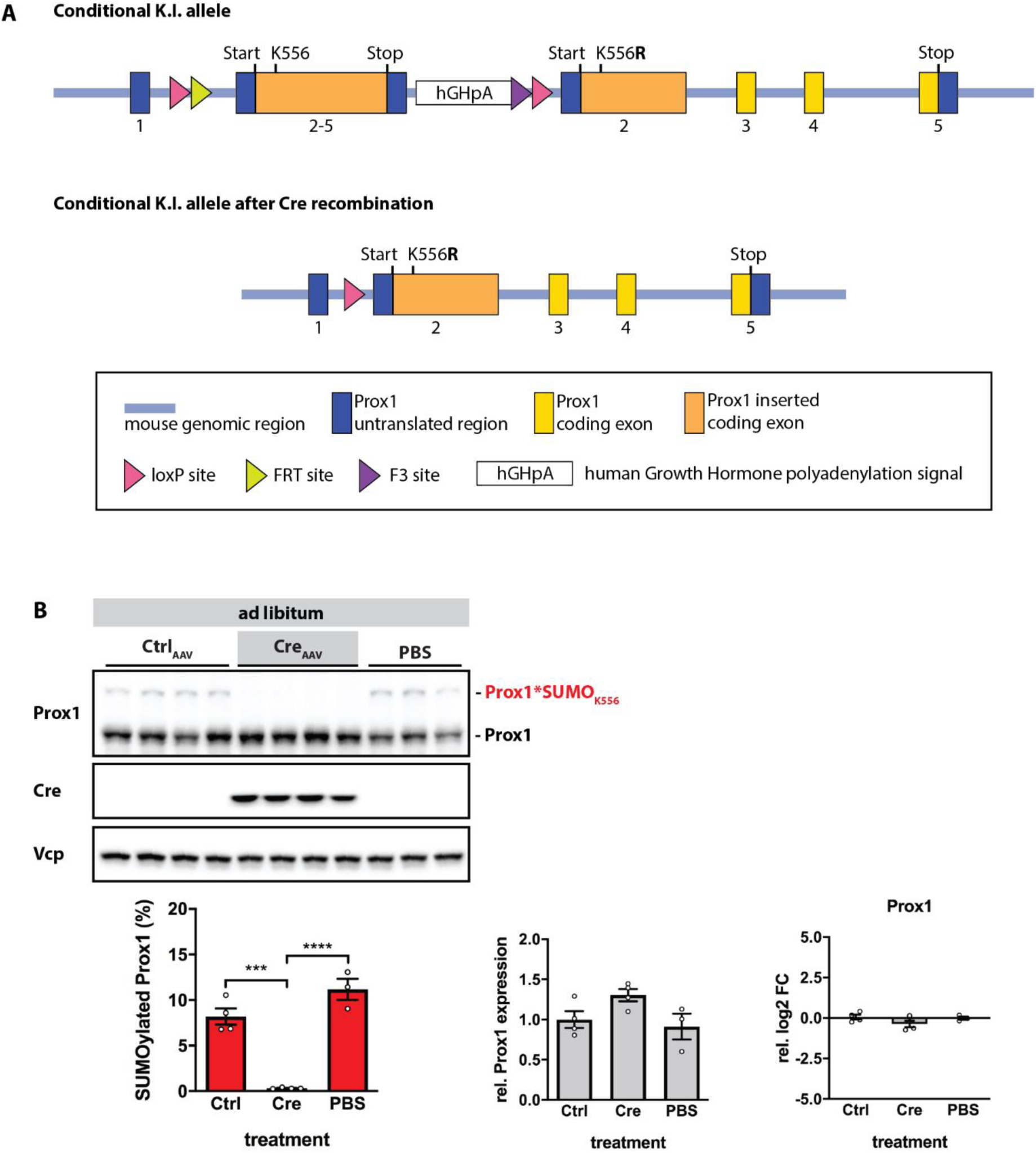
The conditional SUMO-deficient Prox1 knock-in mouse model (Prox1_K556R_ K.I. mice) **A)** Schematic representation of the SUMO-deficient Prox1_K556R_ knock-in mouse model developed together with Taconic Biosciences. **B)** 8 weeks-old Prox1_K556R_ K.I. (f/f) male mice were injected with a control AAV (Ctrl_AAV), with an AAV to overexpress Cre recombinase (Cre_AAV) or with phosphate-buffered saline (PBS) (n=4). Liver tissue samples were collected 3 weeks later from mice fed ad libitum. Liver lysates were analyzed by immunoblotting using anti- Prox1 and anti-Cre antibodies; Vcp was detected for input control. The quantification of SUMOylated Prox1 and total Prox1 protein expression as well as Prox1 mRNA levels analyzed by qPCR are shown. qPCR data presented as relative log2 fold change (FC) normalized to the housekeeping gene TBP. Every dot represents one individual mouse. Data: mean ±SEM. Significance was determined by one-way ANOVA with Tukeýs multiple comparison test between groups. * P≤0.05, ** P≤0.01, *** P≤0.001, **** P≤0.0001.

To demonstrate the validity of our model, eight weeks-old Prox1_K556R_ K.I. male mice were injected with either the Ctrl_AAV, the Cre_AAV, or PBS. The status of Prox1 SUMOylation in the liver was analyzed three weeks later. Although the expression of Prox1 at the mRNA and protein levels was constant between the groups, conjugation of Prox1 with SUMO was abolished in mice injected with the Cre_AAV (Fig. 4B). These results confirm that the main SUMOylation event on hepatic Prox1 happens on lysine 556. The hepatocyte-specificity of the Cre recombination driven by the Cre_AAV was confirmed via liver fractionation experiments (Fig. S2A-B).

We then characterized a cohort of male Prox1_K556R_ K.I. mice injected with PBS, Ctrl_AAV or Cre_AAV fed a standard chow diet for eleven weeks. Blood and tissue samples were collected and examined 3-5 h into the dark phase in the fasted and re-fed states. We identified no differences in terms of body composition, glucose tolerance, insulin sensitivity, lipid metabolism, markers for liver damage, liver weight nor liver morphology (Fig. S3 and S4). In line with the absence of any clear phenotype, the liver transcriptome of the control and the mice expressing the mutant Prox1 was comparable (data not shown). A parallel study of female Prox1_K556R_ K.I. mice fed a standard diet did not reveal differences between the control and the animals expressing the mutant Prox1 (data not shown), overall supporting the notion that the fasting-sensitive SUMOylation switch on Prox1 may only be functionally relevant under dietary stress conditions.

To test this assumption, and given the insensitivity of the Prox1 SUMO-switch under high fat dietary conditions (Fig. 3A) as well as the suggested role of Prox1 in lipid homeostasis, we next characterized a cohort of male Prox1_K556R_ K.I. mice injected with the Ctrl_AAV or Cre_AAV fed a high fat (45%) high fructose (20% w/v) diet for 18 weeks. We selected this diet to induce a metabolic burden at the level of both fatty acid utilization and de novo synthesis. Both groups developed an obese phenotype to a comparable degree, showing a constant increase in body weight and high fasting blood glucose levels (Fig. S5A). At the end of the study, blood and liver samples were collected and examined 3-5 h into the dark phase in the fasted and re-fed states. As expected, the conjugation of Prox1 with SUMO was abolished in mice injected with the Cre_AAV (Fig. 5A). In control animals, the SUMOylated species of Prox1 was detected in both fasted and re-fed states, showing again the insensitivity of the Prox1 SUMO-switch to fasting cues (Fig. 5A).

**Fig. 5.**
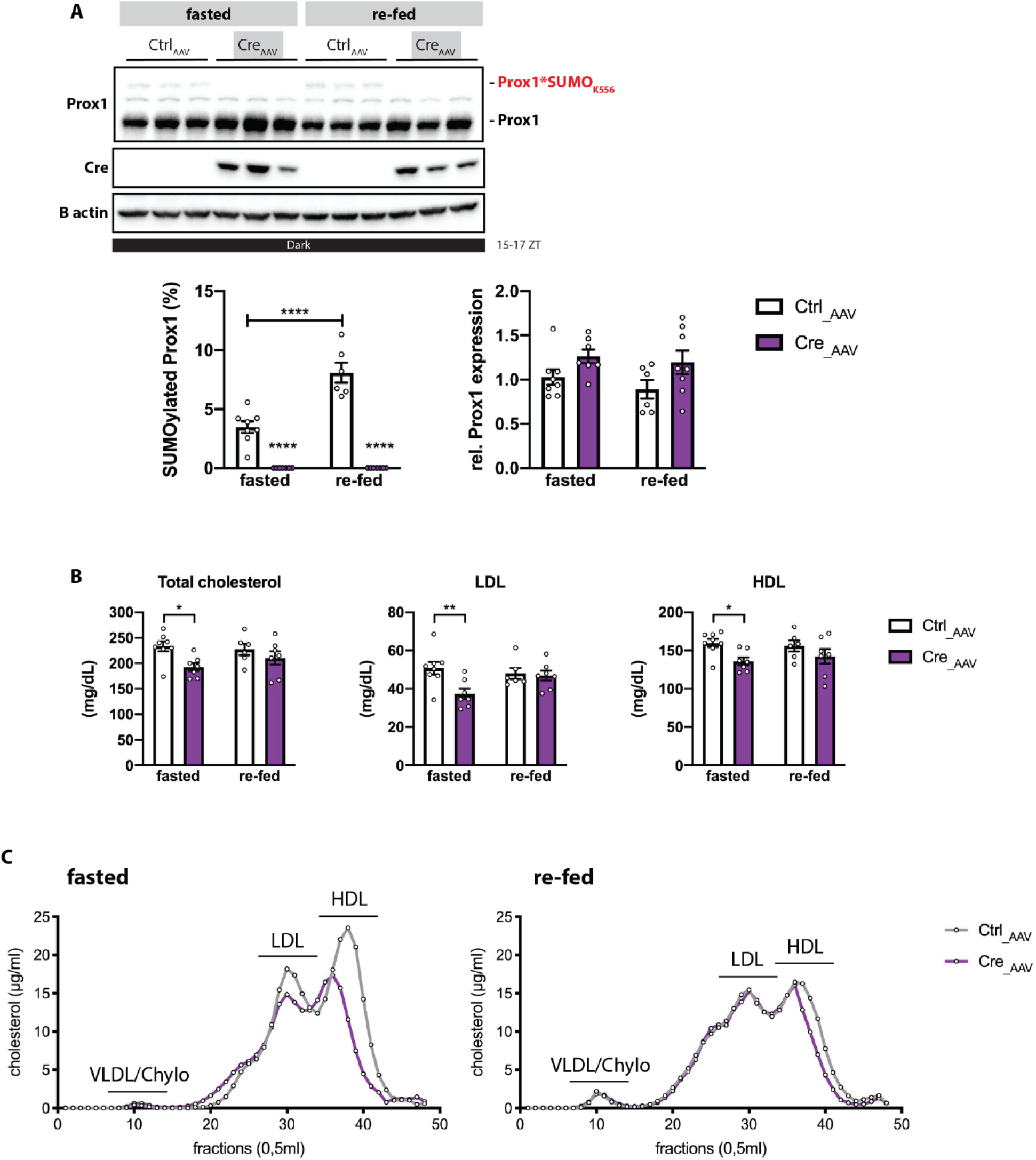
Obese mice expressing SUMO-deficient Prox1 in the liver show reduced serum cholesterol. **A)** 8 weeks-old Prox1_K556R_ K.I. (f/f) male mice were injected with a control AAV (Ctrl_AAV) or with an AAV to overexpress Cre recombinase (Cre_AAV) and placed on a high-fat (45%) high-fructose (20% w/v) diet for 18 weeks. Fasted: the food was removed at ZT 7, samples were collected 8 h later between ZT 15 and 16. Re-fed: mice were fasted for 8 h (ZT 4-12) for synchronization. The food was re-introduced at ZT 12 right before the start of the dark phase, samples were collected between ZT 16 and 17 (n=7-8). Liver lysates were analyzed by immunoblotting using anti-Prox1 and anti-Cre antibodies; B-actin was detected for input control. The quantification of SUMOylated Prox1 and total Prox1 protein expression are shown. **B)** Serum analysis using a colorimetric-based serum analyzer. Levels of total cholesterol, high density lipoprotein (HDL) and low density lipoprotein (LDL) are shown. **C)** Equal amounts of serum were pooled and separated by FPLC, serum lipoprotein cholesterol profiles are shown. A and B) Every dot represents one individual mouse. Data: mean ±SEM. Significance was determined by two-way ANOVA with Sidak’s multiple comparison test between different groups and conditions. * P≤0.05, ** P≤0.01, *** P≤0.001, **** P≤0.0001.

Of note, we identified significantly lower levels of total cholesterol in the circulation of mice expressing the mutant Prox1 as compared to control mice in the fasted state; the cholesterol reduction was reflected in lower levels of low-density lipoprotein (LDL) and high-density lipoprotein (HDL) associated cholesterol (Fig. 5B). No differences in serum triglycerides, bile acids nor markers for liver damage between the control and the mice expressing the mutant Prox1 were identified (Fig. S5A-B).

To corroborate the cholesterol phenotype we performed a serum lipoprotein profile by fast protein liquid chromatography (FPLC). In both groups, we measured high levels of LDL and HDL cholesterol as expected from the high fat/high fructose diet. However, the levels of LDL and HDL associated cholesterol were lower in mice expressing the mutant Prox1 in the fasted state (Fig. 5C). Overall demonstrating that the Prox1 SUMOylation at lysine 556 was specifically responsible for the control of cholesterol handling while leaving other metabolic parameters intact.

We then analyzed the content of cholesterol within the liver and identified no differences between the control and the mice expressing the mutant Prox1 in terms of total cholesterol or cholesteryl esters, which are the storage form of cholesterol in lipoproteins (Fig. S5C). The liver weight and histological parameters were also comparable between the control and the mice expressing the mutant Prox1 (Fig S5C and data not shown).

### Prox1 SUMOylation on lysine 556 controls the hepatic bile acid detoxification pathway

To investigate how the Prox1 SUMO-switch mediated its impact on cholesterol homeostasis under conditions of diet-induced obesity, we compared the liver transcriptome between control and mice expressing the mutant Prox1 by RNA sequencing. A total of 692 differentially expressed transcripts (p < 0.05) were identified in the fasted state, while no transcripts were differentially expressed in the re-fed state (Fig. 6A), demonstrating that the single SUMOylation on lysine 556 controls a distinct subset of downstream target genes during fasting.

**Fig. 6.**
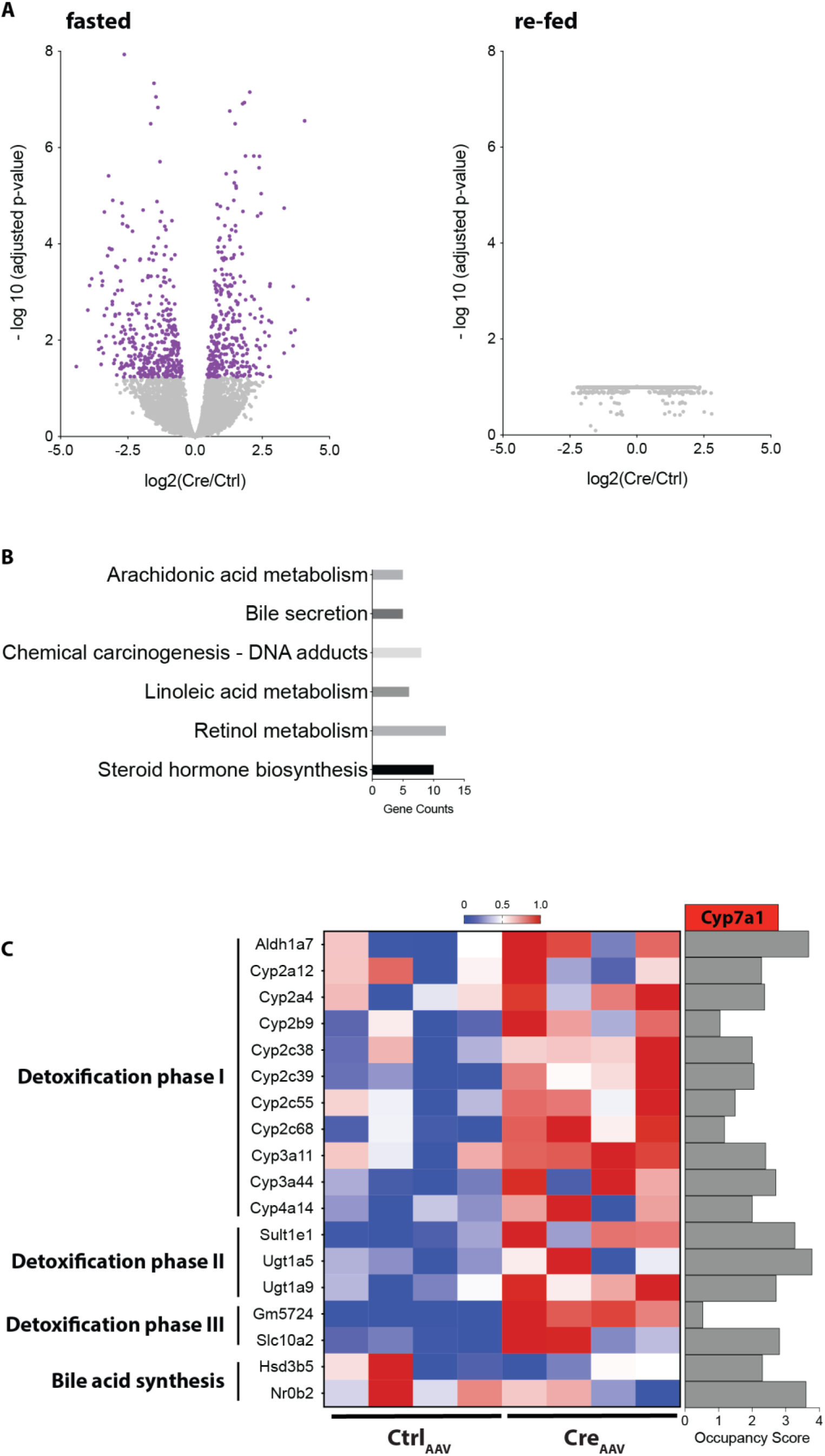
The SUMO-switch on Prox1 controls the hepatic bile acid detoxification pathway. **A)** Volcano plots illustrating the genes that are significantly up- or downregulated in the liver of obese mice expressing the Prox1 K556R mutant in comparison to obese mice expressing wild- type Prox1 in the fasted and re-fed states (n=7-8). **B)** Results of a pathway enrichment analysis (KEGG) of the differentially expressed genes in the fasted state. **C)** Heatmap displaying the expression of enriched genes involved in phase I, II and III of bile acid detoxification and bile acid synthesis in the liver of obese mice expressing the Prox1 K556R mutant (Cre_AAV) in comparison to obese mice expressing wild-type Prox1 (Ctrl_AAV) in the fasted state (n=4). The occupation scores according to a published Chip- sequencing analysis on Prox1 (Armour et al., 2017) are shown. Cyp7a1 was used as a reference gene.

Functional mapping analysis (KEGG) of the differentially expressed genes in the fasted state revealed an enrichment of pathways involved in retinol metabolism, steroid hormone biosynthesis as well as omega-6 fatty acid metabolism and bile secretion (Fig 6B). We then examined the function of the individual genes enriched by the pathway analysis during fasting and identified that most of these genes are involved in the detoxification process of xenobiotics and endogenous molecules such as bile acids and other steroids. The metabolic pathway of detoxification in the liver is driven by enzymes that catalyze the oxidation, reduction, or hydroxylation of toxic molecules (phase I), conjugation of functional groups such as sulfate and glucuronic acid (phase II) and export via membrane transporters (phase III) (Li & Chiang, 2013; Xu et al., 2005). Interestingly, the expression of key enzymes mediating phase I such as cytochrome P450 enzymes (CYPs) and aldehyde dehydrogenases (ALDHs) as well as phase II such as sulfotransferases (SULTs) and UDP-glucuronosyltransferases (UGTs) were upregulated in the liver of mice expressing the Prox1 mutant during fasting (Fig 6C).

Furthermore, the predicted gene 5724 coding for the sodium-independent organic anion transport protein (Oatp1a4) and the sodium-dependent bile salt transporter (Slc10a2 also ASBT) were also upregulated in the liver of mice expressing the Prox1 mutant (Fig 6C).

The analysis also demonstrated that 3 beta-hydroxysteroid dehydrogenase 5 (Hsd3b5), a steroid reductase, and small heterodimer partner 1 (Shp1 or NR0B2), a key transcriptional regulator of bile acid synthesis, were downregulated in the liver of mice expressing the Prox1 mutant (Fig 6C).

To identify whether these differentially expressed genes were potential direct targets of Prox1, we merged our RNA sequencing data with a published Chip-sequencing analysis on Prox1 (Armour et al., 2017). Indeed, Prox1 was enriched at the promoters of key enzymes mediating phase I, II and III of bile acid detoxification as well as factors regulating bile acid synthesis, the occupancy score of the well described target of Prox1 Cyp7a1 was used as reference (Fig. 6C) (Ouyang et al., 2013; Qin et al., 2004). Because SUMO conjugation did not affect the stability, localization nor binding affinity to chromatin of Prox1 (Fig. S6A and B), we hypothesize that the attachment of SUMO to Prox1 rather influences its interaction with regulatory co-factors thus modulating transcription of corresponding target genes.

These results suggested that obese mice expressing the mutant Prox1 had a higher rate of bile acid-mediated cholesterol detoxification during fasting, thus leading to overall reduced cholesterol levels. Taken together, our results establish a nutrition-dependent SUMO-switch on Prox1 in the mouse liver, demonstrating how a single SUMOylation event can exert functional control over hepatic and systemic lipid metabolism.

## Discussion

Hepatocytes are specialized cells that are able to sense hormonal cues and orchestrate metabolic programs to maintain energy homeostasis. In this study, we have characterized a nutrient sensing mechanism in the liver that influences cholesterol metabolism. Prox1, a key transcriptional regulator of lipid metabolism, is modified by SUMOylation on lysine residue 556 in the liver of ad libitum and re-fed mice but this modification is abolished upon fasting.

Utilizing our conditional SUMO-deficient Prox1_K556R_ K.I. mouse model, we demonstrate that the metabolism of lean mice expressing the SUMOylation-deficient Prox1 mutant in hepatocytes is comparable to lean mice expressing the wild type Prox1. However, the SUMO-switch is not responsive to fasting cues in the liver of obese mice. Prox1 is highly modified by SUMOylation even if mice have no access to food. As a consequence, the “engineered” de-conjugation of Prox1 with SUMO in obese mice has a strong impact on cholesterol metabolism, i.e. obese mice expressing the SUMO-deficient Prox1 mutant show reduced levels of circulating LDL and HDL cholesterol.

The metabolism of cholesterol to bile acids is the main pathway for cholesterol clearance. Bile acids are synthesized in hepatocytes, conjugated with taurine or glycine, secreted into the bile and stored in the gallbladder. After food ingestion, they are released into the intestine where they are metabolized by intestinal bacteria to produce un-conjugated more hydrophobic secondary bile acids that are essential for lipid solubilization and absorption. About 95% of the bile acids are re-absorbed in the intestinal tract and returned to the liver and re-secreted into the bile. In addition to its role in lipid absorption, bile acids act as signaling molecules regulating a number of metabolic pathways (Hofmann, 2007; Staels & Fonseca, 2009). However, elevated concentrations of bile acids are cytotoxic. Thus, adaptive responses have evolved to decrease the pool of toxic bile acids. This is achieved by decreasing bile acid synthesis, promoting the metabolism of more hydrophilic bile acids and increasing their elimination (Chen et al., 2014; Xie et al., 2001; Xu et al., 2005).

Prox1 has been shown to act as a negative regulator of the hepatocyte nuclear factor 4α (HNF4α) and the liver receptor homolog 1 (LRH-1), transcription factors modulating several aspects of lipid metabolism such as bile acid synthesis and reverse cholesterol transport (Armour et al., 2017; Qin et al., 2004; Stein et al., 2014). Interestingly, the interaction and thus repression activity of Prox1 to LRH-1 is promoted by SUMOylation of LHR-1 on lysine residue 289 (Stein et al., 2014). In addition, Prox1 has been identified as a recurrent member of the repression complexes NuRD and NCoR mediating the recruitment of histone modifying enzymes such as the histone demethylase LSD1 and the histone deacetylases HDAC2 and HDAC3 (Armour et al., 2017; Ouyang et al., 2013). Furthermore, Prox1 has been shown to interact and repress the steroid and xenobiotic receptor (SXR) in HepG2 cells (Azuma et al., 2011). SXR is a sensor of potentially harmful compounds that activates the expression of genes involved in the detoxification pathway (Xie et al., 2001).

Obese mice expressing a SUMO-deficient Prox1 mutant in the liver show higher transcription levels of genes involved in bile acid detoxification during fasting. Cyp3a4, a crucial enzyme that catalyzes bile acid hydroxylation (Chen et al., 2014) is highly upregulated in mice expressing the mutant Prox1. Bile acid hydroxylation increases the hydrophilicity of bile acids and thus decreases their toxicity. Hydroxylation of bile acids also facilitates their modification by glucuronidation and sulfation which increases their solubility and promotes transport and detoxification (Alnouti, 2009; Perreault et al., 2018). Mice expressing the mutant Prox1 show higher levels of the glucuronosyltransferases Ugt1a5 and Ugt1a9 as well as sulfotransferase Sult1e1. Furthermore, a number of enzymes from the Cyp2c family are also upregulated in mice expressing the mutant Prox1. Enzymes from the Cyp2c cluster catalyze primary bile acids into the more hydrophilic and thus less toxic muricholic acids; this is a process exclusive to the mouse and rat liver (Oteng et al., 2021).

Affecting the composition of the bile acid pool can have significant consequences on cholesterol metabolism. Transgenic mice lacking the Cyp2c cluster and thus lacking muricholic acids have reduced rates of bile acid synthesis and elevated levels of LDL cholesterol (Oteng et al., 2021; Straniero et al., 2020). On the other hand, promoting the synthesis pathway for muricholic acids results in lower levels of LDL cholesterol and an increase in fecal cholesterol excretion (Bonde et al., 2016). Furthermore, an increase in hydrophilic bile acids reduces cholesterol solubility and absorption in the intestine (Wang et al., 2003).

Mice expressing the SUMO-deficient Prox1 mutant also show increased levels of the predicted gene 5724 coding for the sodium-independent organic anion transport protein Oatp1a4. Oatp1a4 is involved in the uptake of unconjugated bile acids into hepatocytes for further metabolism and elimination (van de Steeg et al., 2010). In addition, mice expressing the mutant Prox1 show higher levels of the sodium dependent bile acid transporter Slc10a2 also called ASBT. ASBT is expressed at the membrane of cholangiocytes, the epithelial cells lining the bile ducts, and mediates the re-absorption of bile acids back to hepatocytes for re-secretion into bile (Alpini et al., 2001; Lazaridis et al., 1997; Xia et al., 2006). The recycling of un-conjugated hydrophobic bile acids has the potential to increase the efficiency of bile acid flow and enhance bile acid and cholesterol excretion; this process is referred to as the cholehepatic shunt pathway (Hofmann, 1989).

Interestingly, most of the genes altered by the expression of the SUMO-deficient Prox1 mutant are targets of SXR, which is activated by the accumulation of hydrophobic acid metabolites (Li & Chiang, 2013). Accumulation of bile acids also activates farnesoid X receptor (FXR), which induces the expression of Shp1. Shp1 interacts and represses LRH-1 to inhibit the expression of the rate limiting enzyme for bile acid synthesis generating a negative feed-back loop (Goodwin et al., 2000). As mentioned before, the ability of LHR-1 to induce the expression of genes involved in the absorption, metabolism and excretion of excessive cholesterol is blocked by its SUMO-dependent interaction with Prox1 (Stein et al., 2014).

Therefore, we propose that upon fasting signals, Prox1 is de-SUMOylated to release its repression activity and promote the transcription of genes involved in bile acid detoxification. This could represent a protective mechanism to avoid toxicity of bile acids that accumulate in the liver and gallbladder when food is not available. Loss of SUMOylation on Prox1 could release its repressive activity on a selected set of nuclear receptors resulting in higher rates of bile acid synthesis, bile acid detoxification and bile acid excretion. This mechanism would become especially relevant under metabolic stress that inflicts pressure on cholesterol metabolism. This hypothesis may explain why the metabolism between lean mice expressing the SUMO-deficient Prox1 and the lean controls is comparable, while obese mice expressing the SUMO-deficient Prox1 mutant have lower LDL and HDL cholesterol levels compared to the obese controls during fasting.

Together, we have identified a molecular switch in the mouse liver regulated by nutrient availability. In this respect, the SUMO-switch on Prox1 represents an example of SUMOylation as a mechanism to fine-tune transcriptional programs in response to dynamic environmental cues. The maintenance of nutrient-sensitive SUMOylation on Prox1 may thus contribute to the development of “fasting-based” approaches towards improved metabolic health in the future.

## Author Contributions

Project conceptualization: Stephan Herzig, Julia Szendrödi and Frauke Melchior Formulation of hypothesis, data generation and analysis: Ana Jimena Alfaro N Preliminary data: Claudia Dittner, Janina Becker

Analysis of Bioinformatic data: Anne Loft, Amit Mhamane

Experimental support: Adriano Maida, Anastasia Georgiadi, Phivos Tsokanos, Katarina Klepac, Eveline Molocea, Rabih Merahbi, Karsten Motzler, Julia Geppert, Rhoda Anane Karikari, Susanna Hofmann

## Acknowledgements

We would like to thank Andrea Takas, Elena Vogl, Jeanette Biebl, Daniela Hass and Sebastian Cucuruz for their technical expertise. We thank Luke Harrison for proof reading, editorial support and the graphical abstract. The graphical abstract and part of figure 1 was created with Biorender.com.

---

**Fig. S1 related to Fig. 1.**
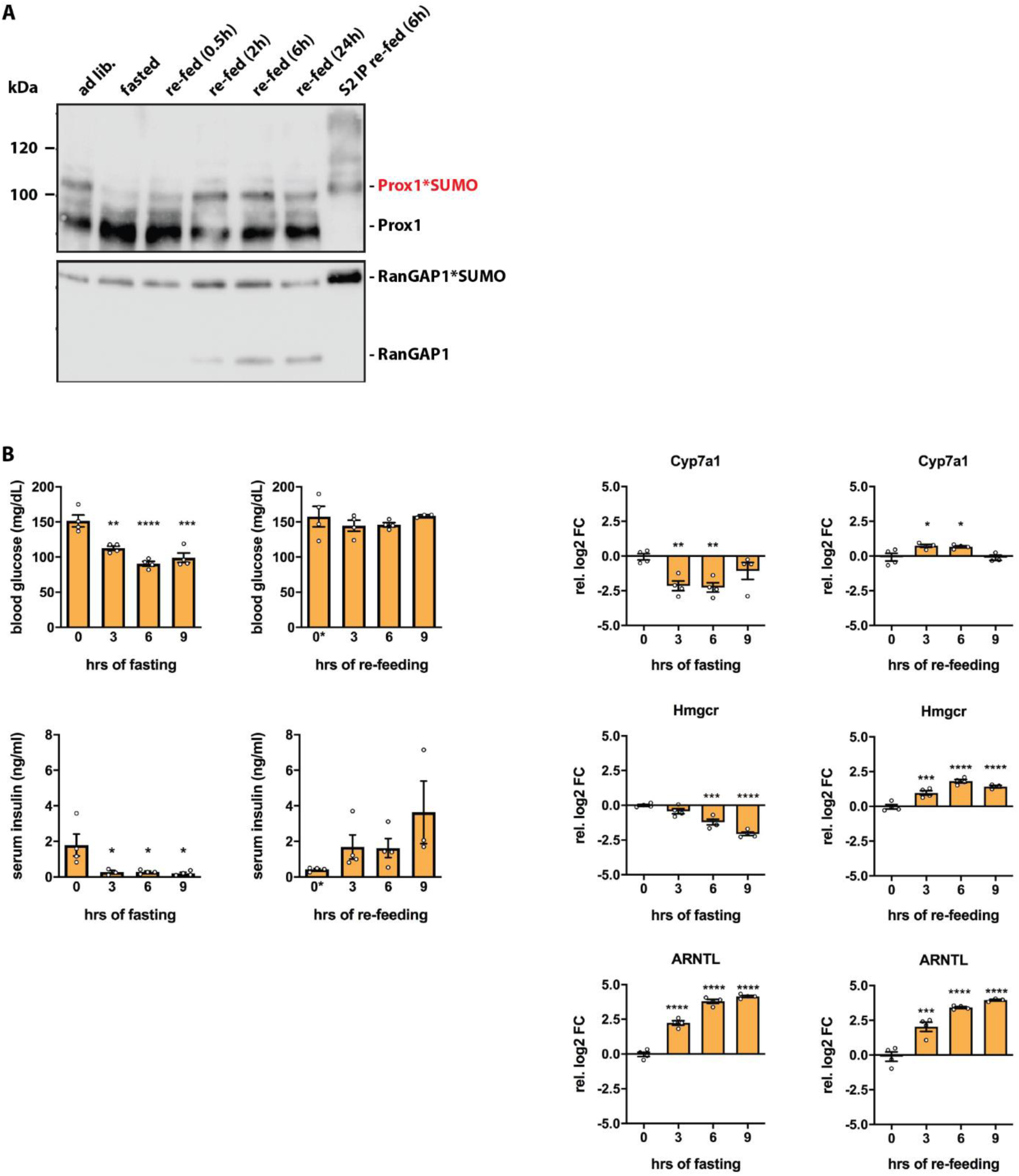
Hepatic Prox1 is modified by SUMOylation in response to fasting-feeding cues. **A)** Liver samples from fasted: (16 h) or re-fed (16 h fasted and re-fed) C57BL/6J mice were analyzed by immunoblotting using anti-Prox1 and RanGAP1 antibodies. The SUMO2 immunoprecipitation eluate was used for size verification. SUMOylated RanGAP1 was used as a loading control. **B)** 8 weeks-old C57BL/6N male mice were fasted: food was removed at ZT 12 or re fed: all groups were fasted for 8 h during the light phase (ZT 4-12) for synchronization, the food was reintroduced at ZT 12. Tissue samples were collected at ZT 12, 15, 18 and 21 (n=4). Blood glucose and insulin levels as well as liver expression analysis at the mRNA level by qPCR are shown. qPCR data presented as relative log2 fold change (FC) normalized to the housekeeping gene TBP. Every dot represents one individual mouse. Data: mean ±SEM. Significance was determined by one-way ANOVA with Dunnett’s multiple comparison test relative to samples collected at ZT 12. * P≤0.05, ** P≤0.01, *** P≤0.001, **** P≤0.0001.

**Fig. S2 related to Fig. 4.**
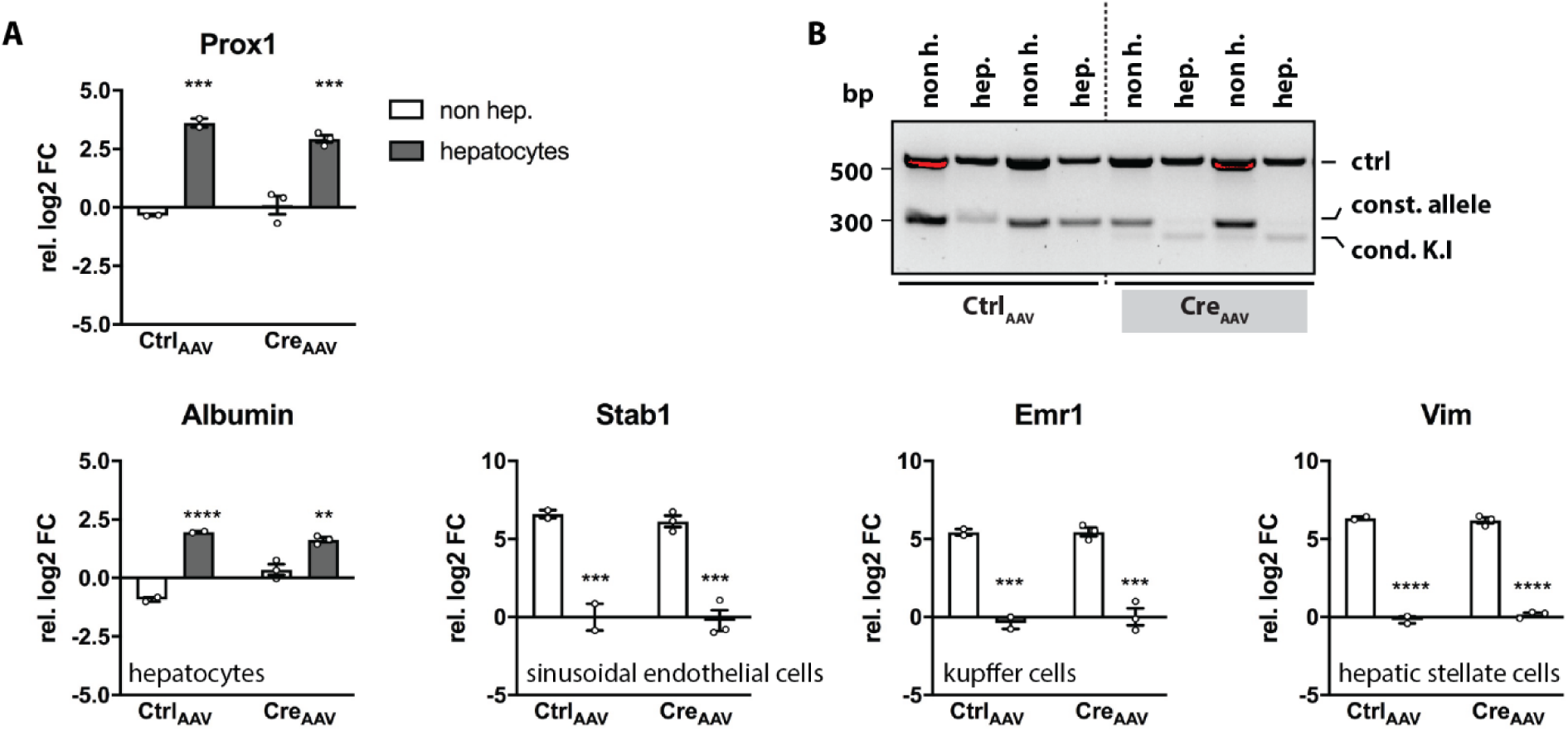
The conditional SUMO-deficient Prox1 knock-in mouse model (Prox1K556R K.I. mice) **A)** 8 weeks-old Prox1_K556R_ K.I. (f/f) male mice were injected with a control AAV (Ctrl_AAV) or an AAV to overexpress Cre recombinase (Cre_AAV) (n=2-4). 3 weeks later hepatocytes (hep.) were isolated from the non-hepatocyte (non.h.) fraction. Expression analysis at the mRNA level by qPCR, data presented as relative log2 fold change (FC) normalized to the housekeeping gene TBP. Every dot represents one individual mouse. Data: mean ±SEM. Significance was determined by two-way ANOVA with Sidak’s multiple comparison test between different conditions. *P≤0.05, ** P≤0.01, *** P≤0.001, **** P≤0.0001. **B)** Amplification reaction by PCR detecting the constitutive allele (280 bp) or the conditional K.I. allele after Cre mediated recombination (235 bp); control product (585 bp).

**Fig. S3.**
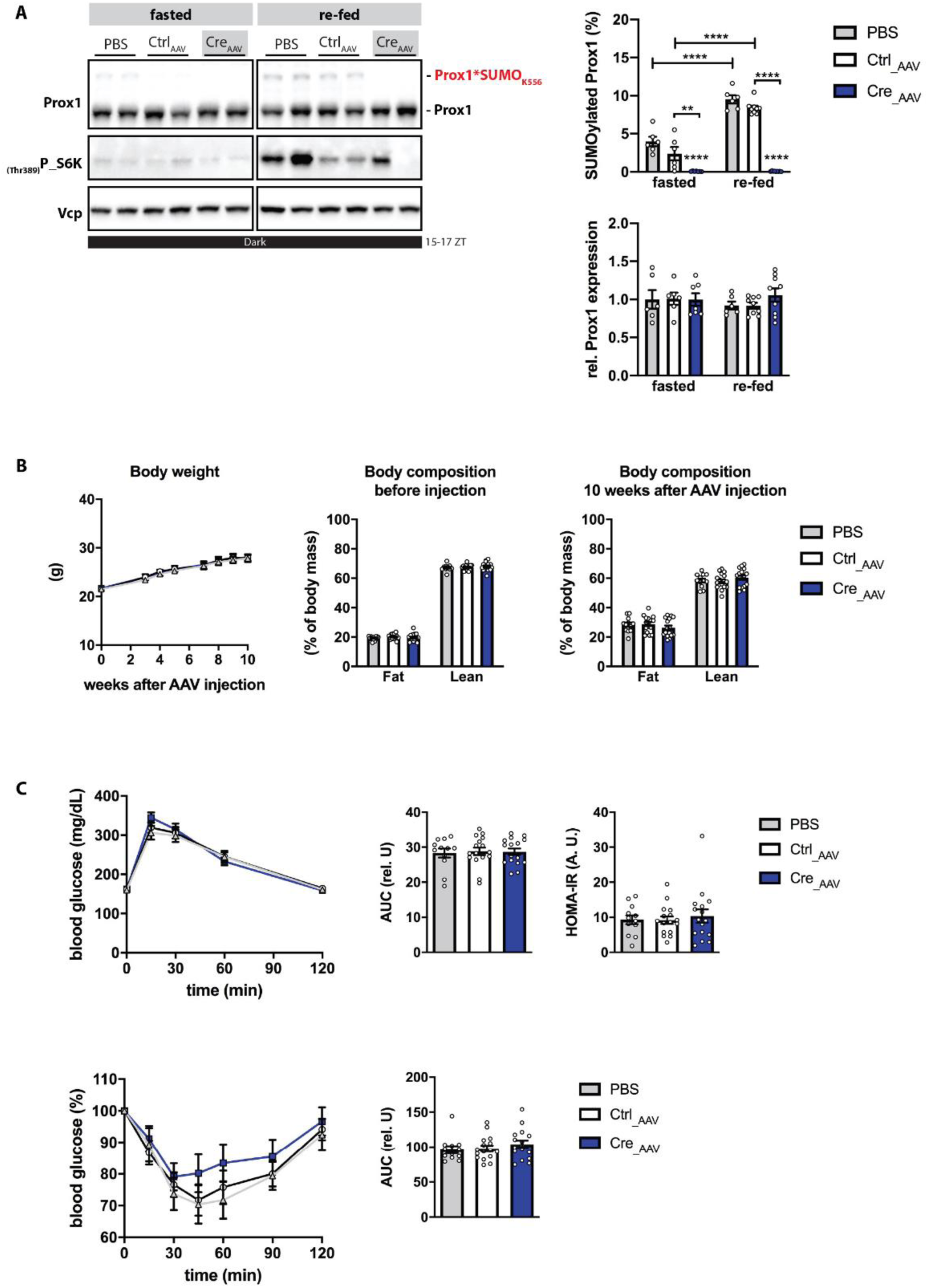
Characterization of male mice expressing a Prox1_K556R_ mutant in the liver (part 1) 8 weeks-old Prox1_K556R_ K.I. (f/f) male mice were injected with phosphate-buffered saline (PBS), a control AAV (Ctrl_AAV) or with an AAV to overexpress Cre recombinase (Cre_AAV). 11 weeks after the AAV injection the study was terminated. Fasted: the food was removed at ZT 7, samples were collected 8 h later between ZT 15 and 16. Re fed: mice were fasted for 8 h (ZT 4- 12) for synchronization, the food was re-introduced at ZT 12 right before the start of the dark phase. Samples were collected between ZT 16 and 17 (n=6-9). **A)** Liver lysates were analyzed by immunoblotting using anti-Prox1 and anti-P_S6K(Thr389) antibodies; Vcp was detected for input control. The quantification of SUMOylated Prox1 and total Prox1 protein expression are shown. Every dot represents one individual mouse. Data: mean ±SEM. Significance was determined by two-way ANOVA with Sidak’s multiple comparison test between different groups and conditions. **B)** Body weight records and Echo-MRI measurements taken before and 10 weeks after the AAV injections are shown (n=12-18). **C)** Top: GTT performed 8 weeks after the AAV injection. For the GTT mice were fasted for 5 to 6 h then challenged with glucose (1.5 g/kg) via an intraperitoneal injection. Blood samples were collected at time points: 0, 15, 30, 60 and 120 min after the glucose injection (n=12-18). Bottom: ITT performed 9 weeks after the AAV injection. For the ITT mice were fasted for 5 h then challenged with insulin (1.2 U/kg) via an intraperitoneal injection. Blood samples were collected at time points: 0, 15, 30, 45, 60, 90 and 120 min after the insulin injection (n=12-18). The area under the curve (AUC) and the calculated homeostatic model assessment for insulin resistance (HOMA-IR) are shown. (B-C) Every dot represents one individual mouse. Data: mean ±SEM. Significance was determined by one-way ANOVA with Tukeýs multiple comparison test between different groups.

**Fig. S4.**
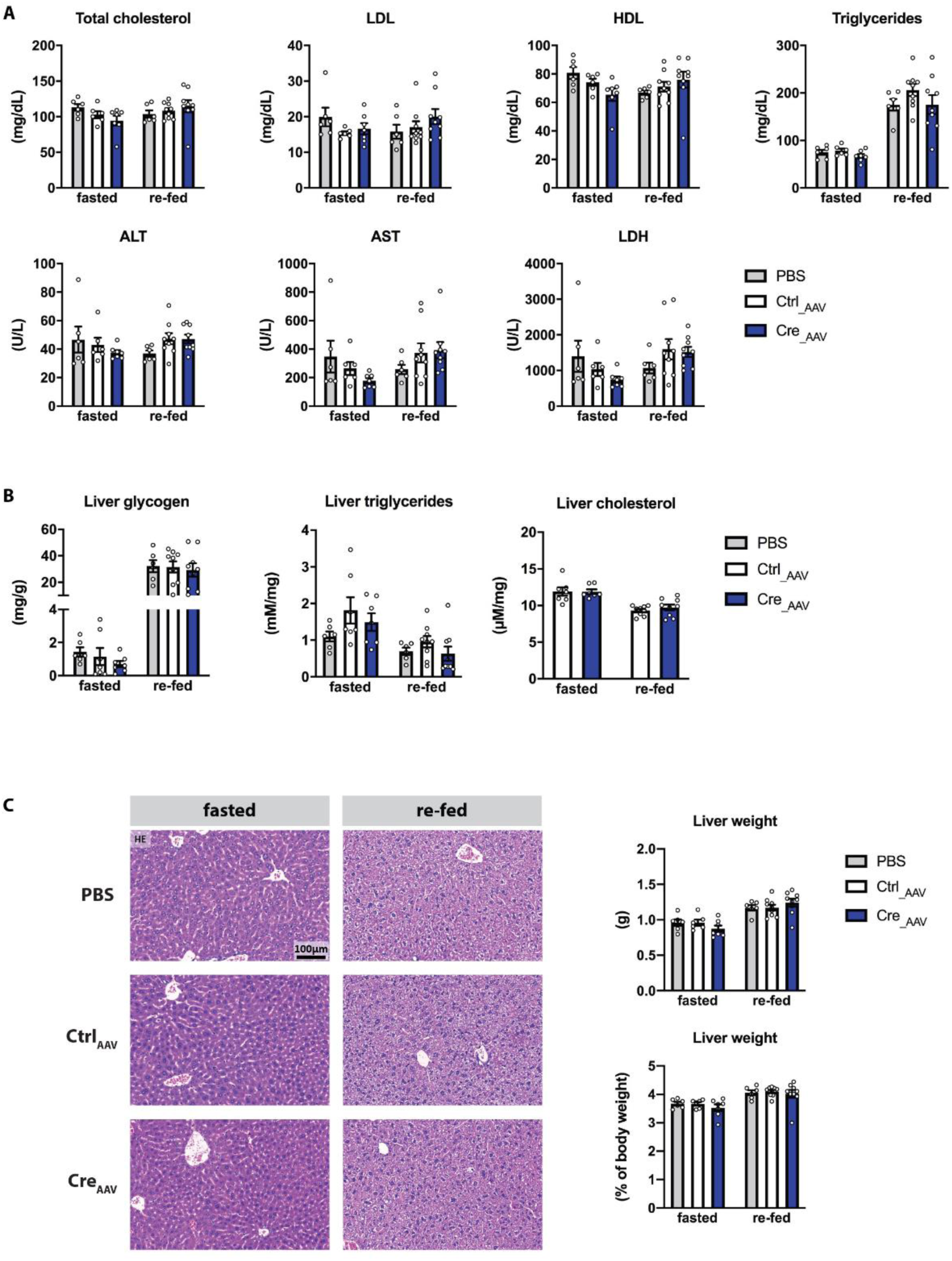
Characterization of male mice expressing a Prox1_K556R_ mutant in the liver (part 2) 8 weeks-old Prox1_K556R_ K.I. (f/f) male mice were injected with phosphate-buffered saline (PBS), a control AAV (Ctrl_AAV) or with an AAV to overexpress Cre recombinase (Cre_AAV). 11 weeks after the AAV injection the study was terminated. Fasted: the food was removed at ZT 7, samples were collected 8 h later between ZT 15 and 16. Re fed: mice were fasted for 8 h (ZT 4- 12) for synchronization, the food was re-introduced at ZT 12 right before the start of the dark phase. Samples were collected between ZT 16 and 17 (n=6-9). **A)** Serum analysis using a colorimetric-based serum analyzer. Levels of total cholesterol, low density lipoprotein (LDL), high density lipoprotein (HDL), triglycerides, alanine aminotransferase (ALT), aspartate aminotransferase (AST) and lactate dehydrogenase (LDH) are shown. **B)** Colorimetric measurement of glycogen, triglycerides and total cholesterol content within the liver. **C) Left:** Liver morphology analyzed via hematoxylin and eosin staining, representative images. **C) Right:** Liver weight (in grams and % relative to the total body weight). Every dot represents one individual mouse. Data: mean ±SEM. Significance was determined by two-way ANOVA with Sidak’s multiple comparison test between different groups and conditions.

**Fig. S5 related to Fig. 5.**
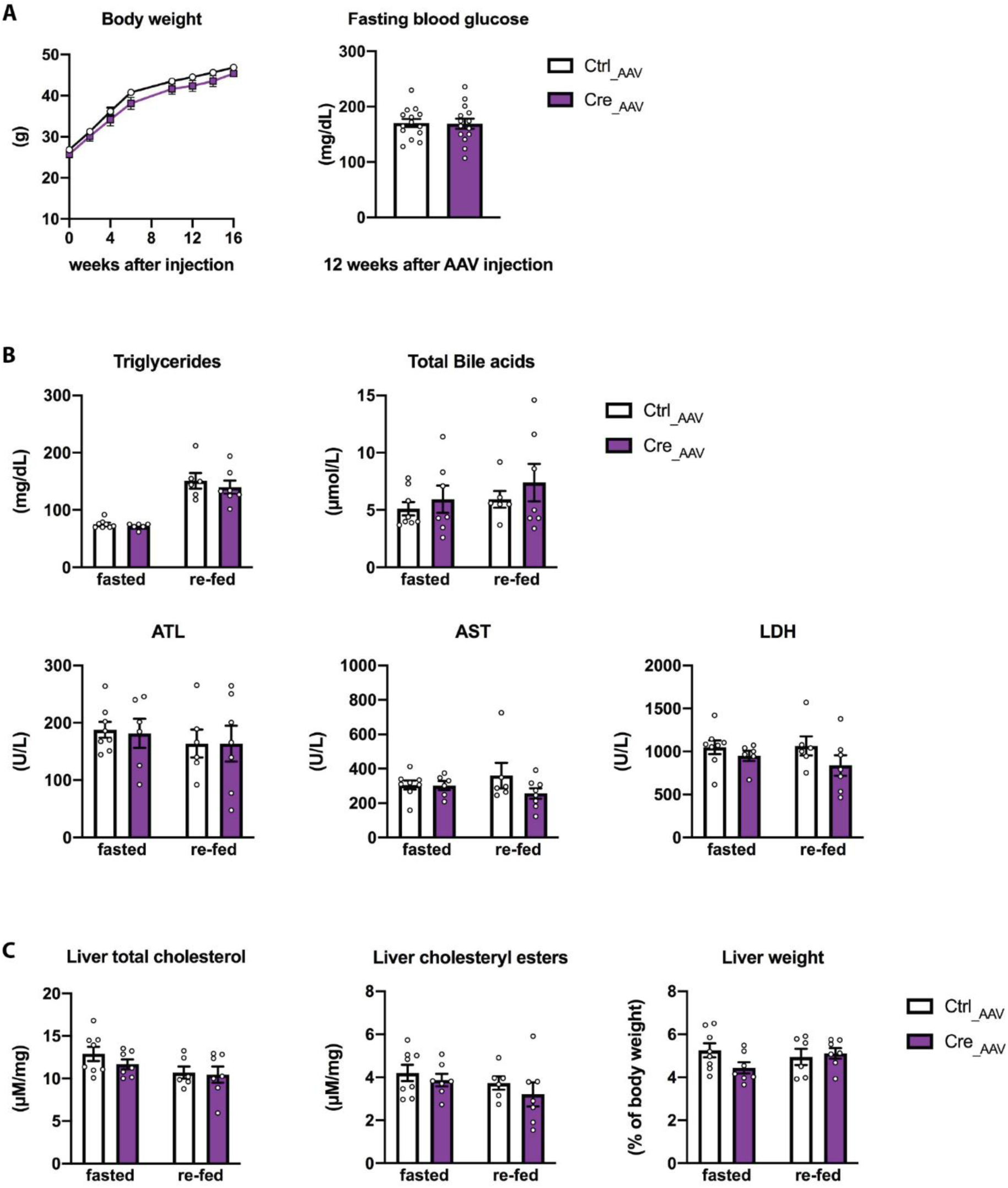
The SUMO-switch on Prox1 regulates cholesterol metabolism in obese mice. 8 weeks-old Prox1K556R K.I. (f/f) male mice were injected with a control AAV (Ctrl_AAV) or with an AAV to overexpress Cre recombinase (Cre_AAV) and placed on a high-fat (45%) high-fructose (20% w/v) diet for 18 weeks. Fasted: the food was removed at ZT 7, samples were collected 8 h later between ZT 15 and 16. Re-fed: mice were fasted for 8 h (ZT 4-12) for synchronization. The food was re-introduced at ZT 12 right before the start of the dark phase, samples were collected between ZT 16 and 17 (n=7-8). **A)** Body weight records and fasting glucose levels (recorded 12 weeks after the AAV injection) are shown (n=14). Every dot represents one individual mouse. Data: mean ±SEM. Significance was determined by two-tailed, unpaired t-test. **B)** Serum analysis using a colorimetric-based serum analyzer. Levels of triglycerides, total bile acids, alanine aminotransferase (ALT), aspartate aminotransferase (AST) and lactate dehydrogenase (LDH) are shown (n=7-8). **C)** Colorimetric measurement of total cholesterol and cholesteryl ester within the liver as well as liver weight (in % relative to the total body weight) are shown (n=7-8). Every dot represents one individual mouse. Data: mean ±SEM. Significance was determined by two-way ANOVA with Sidak’s multiple comparison test between different groups and conditions. *P≤0.05, ** P≤0.01, *** P≤0.001, **** P≤0.0001.

**Fig. S6.**
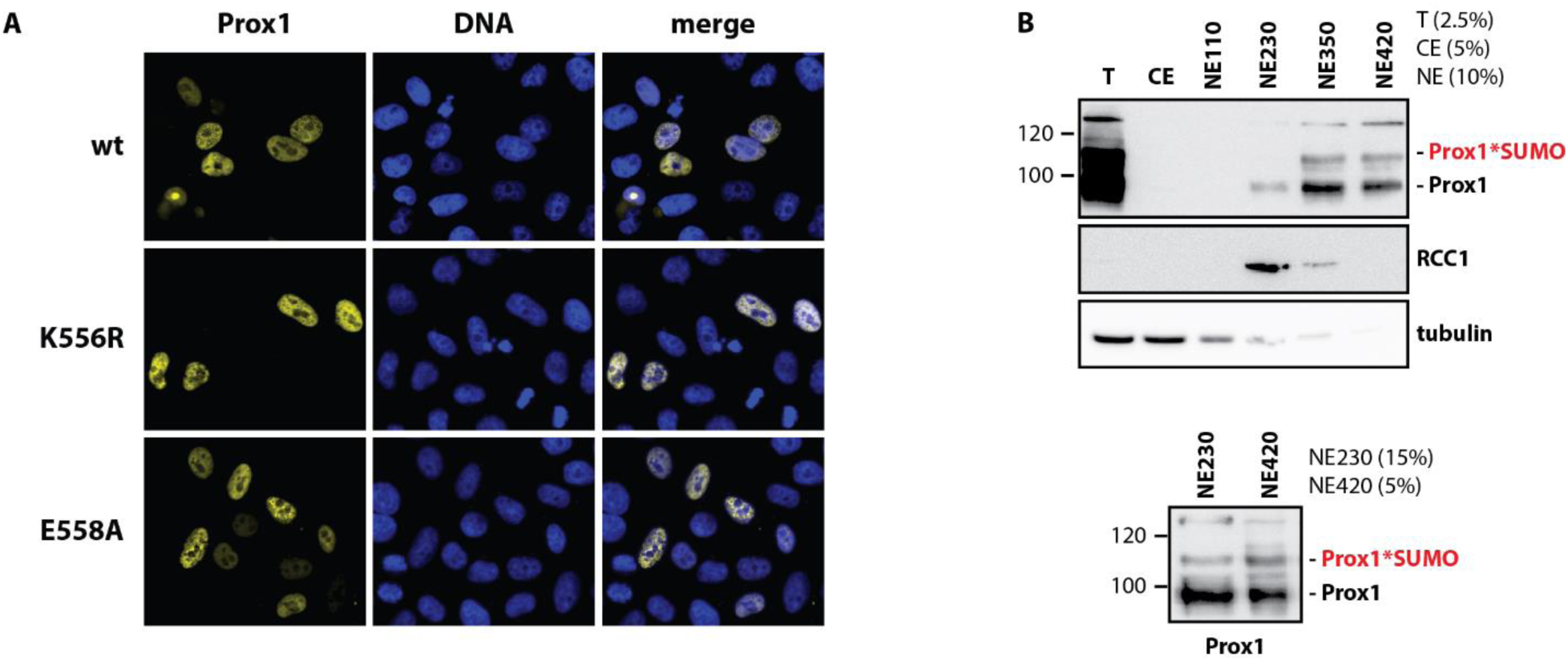
Prox1 SUMOylation does not affect the stability, localization nor binding affinity to chromatin. **A)** HeLa cells were transfected with mouse HA-tagged wild-type Prox1 (wt), K556R mutant Prox1 (KR) or E558A mutant Prox1 (EA). Localization of Prox1 was analyzed by immunofluorescence using anti-HA antibodies (yellow), DNA was stained with Hoechst (blue). **B)** The nuclei from HepG2 cells were isolated and used for protein extraction. Multiple nuclear extracts were prepared using different concentrations of NaCl (110, 230, 350 or 420 mM). Samples collected before cell lysis (T), the cytoplasmic extracts (C) and the nuclear extracts (NE) NEs were analyzed by immunoblotting using anti-Prox1 antibodies. The chromatin binding factor RCC1 and tubulin were used as controls.

## Supplemental Experimental Procedures

### Generation of AAV constructs

The pdsAAV_LP1_GFPmut-miNC vector was digested with KnpI and NotI to remove the GFPmut-miNC control sequence. The sequence coding for Cre recombinase was amplified from a bacterial expression construct from St. Judes Children’s Research Hospital, Memphis, TN, USA. Primers were designed to introduce a KnpI restriction site before the Cre recombinase sequence and a NotI restriction site directly after. Parallel amplification reactions were performed to introduce two point mutations and generate two stop codons right after the Kozak sequence. After ligation and endotoxin-free plasmid purification, the LP1_Cre_wt_ and LP1_Cremut sequence integrity and orientation were confirmed by sequencing. To corroborate the integrity of the inverted terminal repeats (ITRs) the plasmids were subjected to a test digestion and the band pattern was controlled by comparison with calculated results. The AAV_LP1_Cre_wt_ (Cre_AAV) and AAV_LP1_Cre_mut_ (Ctrl_AAV) constructs were used for a large scale AAV stereotype 8 packaging and purification done by Vigene Biosciences Rockville, MD, USA; pDGdelta-helper and p5E18-RC plasmids were provided by our laboratory.

### Cre recombination specificity by liver fractionation

8 weeks-old Prox1_K556R_ K.I. (f/f) mice were injected with either the Ctrl_AAV or the Cre_AAV. 3 weeks later the hepatocyte cell fraction was separated from the non-hepatocyte fraction following the protocol generated by (Godoy et al., 2013). In brief: Mice were anaesthetized, both abdominal walls were opened and the liver was perfused through the venae cavae with a warm EGTA-containing KH/HEPES buffer for 10 min. The liver was then perfused with a warm collagenase-KH/HEPES buffer for 12 min until liver digestion was visible. The perfused liver was then removed, incubated in suspension buffer (KH/HEPES containing 400 mg BSA) and dissociated by gentle shaking. The cell suspension was filtered through a 100 nm pore mesh and centrifuged at 50 g for 5 min at 4° C. The pellet containing the hepatocyte fraction was washed 2 times with suspension buffer and centrifuged at 50 g for 5 min at 4° C. The supernatant containing the non-hepatocyte fraction was transferred to a clean container and cleared by a second centrifugation round at 50 g for 5 min at 4° C. The clean supernatant was then centrifuged at 800 g for 10 min to pellet the non-hepatocyte cell fraction. The liver fractionation was controlled by detecting the expression of albumin, an hepatocyte-specific marker, as well as Stab1, Emr1 and Vim, which are markers of sinusoidal endothelial cells, kupffer cells and hepatic stellate cells, respectively.

A PCR was then performed using primers designed by Taconic Biosciences to detect either the constitutive allele or the conditional knock-in (K.I.) allele after Cre-mediated recombination.

Amplification with these primers gives a product of 280 bp for the constitutive allele and a 235 bp for the conditional K.I. allele.

### In vivo characterization protocols

#### Glucose and Insulin tolerance tests

For the glucose tolerance test, mice were fasted for 5 to 6 h (ZT 2-8) then challenged with 1.5 g/kg glucose via an intraperitoneal injection. Blood samples were collected in heparin coated tubes at time points: 0 and 15 min from the tail vein; the blood glucose levels were recorded with a glucometer at time points: 0, 15, 30, 60 and 120 min. Blood samples were centrifuged at 2000 g for 5 min at 4° C; the plasma was collected, snap-frozen in liquid nitrogen and stored at - 80° C. The plasma insulin content was measured with a mouse insulin ELISA (ALPCO - 80-INSMS-E10). For the insulin tolerance test, mice were fasted for 5 h (ZT 2-7) then challenged with 1.2 U/kg insulin via an intraperitoneal injection. Blood glucose levels were recorded with a glucometer at time points: 0, 15, 30, 60 and 120 min from the tail vein.

#### Blood sampling

Blood samples of 20-60 μl volume were collected in heparin coated tubes by piercing the tail vein. Blood samples were centrifuged at 2000 g for 5 min at 4° C; the plasma was collected, snap-frozen in liquid nitrogen and stored at -80° C.

### Histology

Tissue samples were harvested and immediately fixed with neutrally buffered formalin (4 % w/v) (Sigma-Aldrich - HT501128) and subsequently routinely embedded in paraffin (Tissue Tec VIP.6 Sakura Europe). Sections of 3 µm were stained with hematoxylin and eosin (HE), using a HistoCore SPECTRA ST automated slide stainer (Leica) with prefabricated staining reagents (Histocore Spectra H&E Stain System S1 - Leica), according to the manufacturer’s instructions. The stained tissue sections were scanned with an AxioScan.Z1 digital slide scanner (Zeiss) equipped with a 20x magnification objective.

### Liver glycogen, triglycerides and cholesterol measurements

For glycogen measurements: Liver tissue pieces were weighted (45-55 mg) and homogenized in 0.5 ml of a KOH (30 %) solution. Samples were mixed at 1000 rpm for 1 hr at 95° C then centrifuged at 500 g for 5 min at room temperature. The supernatant was collected and mixed with 1.4 ml ice-cold ethanol (95 %), incubated for 30 min at -20° C and centrifuged at 3000 g for 20 min at room temperature. The pellets were washed with ethanol (95 %), dried for 10 min at 60° C and dissolved in 0.25 ml distilled water at 37° C. Glycogen content was measured using a glycogen assay kit (Sigma-Aldrichh) according to the manufacturer’s protocol.

For lipid measurements: Liver tissue pieces were weighted (60-80 mg) and homogenized in 1.5 ml of a chloroform:methanol (2:1) solution. Samples were mixed at 1400 rpm for 20 min at room temperature then centrifuged at 13000 rpm for 30 min at room temperature. The liquid phase was collected and mixed with 0.2 ml NaCl (150 mM) and centrifuged at 2000 rpm for 5 min. 0.2 ml of the organic phase was mixed with 40 μl of a chloroform:Triton-X (1:1) solution and dried overnight with the speed-vac V-AL program in the Concentrate plus (Eppendorf). The Triton-X lipid solution was diluted (1.125x) in 0.2 ml distilled water by mixing (end-to-end rotation) for 1 hr at room temperature. Triglycerides were measured using a Triglycerides determination kit (Sigma-Aldrichh) according to the manufacturer’s protocol using 2 μl of lipid solution. Cholesterol levels were measured using a total cholesterol assay kit (Cell Biolabs INC.) according to the manufacturer’s protocol using 2 μl of lipid solution.

### Purification of full length mouse Prox1

The sequence coding for mouse wild type, K556R or E558A Prox1 were cloned from a pcDNA3 HA-Prox1 into a pEYFP-C1 vector. Primers were designed to introduce a Sal I restriction site followed by a PreScission protease recognition site directly before the translation start site, Xba I restriction site was introduced directly after the translation termination site. Sequence integrity and orientation was confirmed by sequencing.

YFP-Prox1 was transfected into HEK293T cells with polyethylenimine for 48 h. Then the cells were collected with ice-cold PBS and centrifuged at 1000 g for 5 min at 4° C, the cell pellet was washed one more time with ice-cold PBS. Cells were then lysed with 1 ml ice-cold assay buffer containing Tris (50 mM) pH 7.5, EDTA (5 mM), EGTA (5 mM), NaCl (150 mM), Igepal (0.5 %) supplemented with protease inhibitors. Cell lysates were sonified at a 20 % amplitude for 15 pulses (1 sec pulse and 1 sec break), incubated on ice for 30 min then clarified by centrifugation at 20000 g for 30 min at 4° C. GFP-binder beads pre-equilibrated in assay buffer were incubated with the lysates (around 5 μl GFP-binder beads per 1 mg protein) overnight with a constant agitation at 4° C. The GFP-binder beads were collected by centrifugation at 800 g for 5 min at 4°C using a swing-out rotor. The beads were washed twice with 1 ml ice-cold RIPA buffer and three times more with 1 ml ice-cold assay buffer supplemented with DTT (1 mM). During the last wash, the beads were transfer into a 0.5 ml tube in a 300 μl volume assay buffer supplemented with DTT (1 mM) and 2 μg GST-tagged PreScission protease were added. The cleavage was performed for 4 h with a constant agitation at 4° C. The GFP-beads were collected by centrifugation at 800 g for 5 min at 4°C using a swing-out rotor and the supernatant was transfer to a fresh 0.5 ml tube. To remove the GST-tagged PreScission protease, the eluate was incubated with 10 μl GST-binder beads pre-equilibrated in assay buffer supplemented with DTT (1 mM) for 1.5 h with a constant agitation at 4° C. The GST-binder beads were collected by centrifugation at 500 g for 5 min at 4°C using a swing-out rotor and the eluate was concentrated to a 50-100 μl volume using a VIVASPIN 0.5 ml centrifugal concentrator (Millipore) according to the manufacturer’s instructions. The eluate was snap-frozen in liquid nitrogen and stored at -80° C.

### Immunofluorescence Microscopy

HeLa cells were transfected with mouse HA-tagged wild-type Prox1 (wt), K556R mutant Prox1 or E558A mutant Prox1 for 24 h and fixed with 4% formaldehyde in PBS for 20 min at room temperature. Cells were washed with PBS and permeabilized with Triton X-100 (0.2 %) in PBS for 10 min. Cells were blocked with BSA (2 %) in PBS for 1 hr and incubated with anti-HA antibodies (Covance - MMS-101P) for 1 hr in a humid chamber. Cells were washed twice with PBST and once with PBS then incubated with a mouse-Alexa488 antibody and Hoechst (0.2 μg/μl) for 1 hr in the dark. Samples were mounted with Daco and analyzed with an Axioskop2 fluorescence microscope (Zeiss) using a 40x oil immersion objective (NA:1.4).

### Salt-gradient Prox1 extraction from HepG2 cells nuclei

Cells were washed twice with ice-cold PBS by centrifugation at 1000 g for 5 min at 4° C. Cells were then incubated for 3min with 5x pellet volume ice-cold cytoplasmic buffer containing HEPES (10 mM), EDTA (1 mM), EGTA (1mM), KCl (60mM), Igepal (0.075 %) supplemented with protease and phosphatase inhibitors (pH = 7.6). Cell lysis was controlled by trypan blue staining (up to 70-80 % cell lysis). The sample was then centrifuged at 1500 g for 4 min at 4°C and the cytoplasmic extract was transferred to a fresh tube. The nuclear pellet was gently washed with 3x pellet volume ice-cold cytoplasmic buffer by pipetting up and down and centrifuged at 1500 g for 4 min at 4° C. The nuclear pellet was then incubated for 10 min with 2x pellet volume ice-cold nuclear buffer containing HEPES (20 mM), MgCl_2_ (1.5 mM), EGTA (0.2 mM), glycerol (25 %) supplemented with protease and phosphatase inhibitors (pH = 7.9). Parallel extractions were done using the nuclear buffer supplemented with different NaCl concentrations (110, 230, 350 or 420 mM). Samples were vortexed periodically to re-suspend the pellet. Finally, the nuclear extracts were centrifuged at 1500 g for 4 min at 4°C and transferred to a fresh tube. The cytoplasmic and nuclear extracts were clarified by centrifugation at 20000 g for 30 min at 4° C. Samples were snap-frozen in liquid nitrogen and stored at -80° C.

